# Cellular-resolution Three-photon Ultra-Deep Imaging (TrUDI) of neuronal population activity at 2 mm depth in the mammalian brain

**DOI:** 10.64898/2026.10.16.759624

**Authors:** Kalyan Santra, Francesca Logiacco, Agustina Frechou, Kevin Barber, Arash Bellafard, Jyun-you Liou, Alipasha Vaziri

## Abstract

Extending the depth reach of multi-photon microscopy for imaging neuronal population activity in the scattering mammalian brain is an “exponentially hard” problem. Three-photon (3p) microscopy has enabled imaging of neuroactivity at depths up to ∼1.2 mm in the mouse brain. However, the fundamental depth limit of 3p imaging, as set by the depth at which the out-of-focus background fluorescence exceeds the in-focus fluorescence signal, is well beyond 2 mm. Reaching this fundamental depth limit in 3p imaging has been hampered by insufficient in-focus fluorescence signal arising from suboptimal optical design and acquisition conditions. Here, we demonstrate a Three-photon Ultra-Deep Imaging (TrUDI) system that approaches this fundamental depth limit via systematic and principled optimization of the in-focus fluorescence signal. TrUDI optimally matches the parameters of the excitation laser, the optical system, and the acquisition to linear and non-linear limits for safe laser exposure. At the same time, TrUDI remains compatible with the spatial and temporal requirements for single-cell resolution calcium imaging, and its parameters are tuned to the photo-physics of 3p excitation and the scattering properties of the mouse brain tissue. Our approach results in a ∼250-fold increase in detected fluorescence photons at the same laser power as in previous approaches. We use TrUDI to demonstrate *in vivo* cellular-resolution recordings of neuronal population activity within fields of view of ∼230 × 230 µm^2^ at 8.6 Hz at tissue depths of up to ∼2 mm in mouse cortex and hippocampus.

## Main

The functional complexity of the mammalian brain arises from its highly interconnected network across regions and depths, each performing specialized roles such as integrating sensory information^1–3^, executing parallel computations^4–7^, and coordinating behavioral responses^8–16^. These functions are distributed across brain-wide neuronal populations, which communicate through dense, long-range connections^17–20^. In recent decades, the combination of various two-photon fluorescence microscopy (2pM) techniques^21–25^ and genetically encoded calcium indicators (GECIs) such as GCaMP^26–30^ has enabled large-scale recording of neuronal population dynamics at the cellular level in scattering mammalian brain. However, imaging the activity of neurons in sub-cortical regions such as the hippocampus, the thalamus or the striatum has remained highly challenging.

The nonlinearity of excitation in 2pM leads to spatial confinement of the fluorescence signal and suppression of out-of-focus fluorescence and thus allows to resolve cellular and sub-cellular features within the depth range of the mouse cortex. However, the generated fluorescence signal in 2pM degrades exponentially as a function of tissue depth due to scattering and absorption loss of excitation photons, as characterized by the *effective attenuation length* (EAL) of the tissue^31,32^. This imposes a fundamental limit on the depth reach of 2pM^33–35^, as this loss cannot be arbitrarily compensated by an increase in the excitation power. At some critical depth (∼600–700 µm in the mouse cortex^36,37^) the resulting energy density of the excitation light at the tissue surface will result in a background fluorescence that exceeds the generated signal at the focus, resulting in unity signal-to-background ratio (SBR).

In this context, three-photon fluorescence microscopy (3pM)^32,37–41^ has emerged as a promising solution. The advantages of 3pM stem from two factors: First, the higher-order nonlinearity of excitation in 3pM leads to a significantly lower out-of-focus fluorescence background at the same tissue depth compared to 2pM, and as a result pushes the limit of unity SBR to much greater tissue depths. Second, scattering at the longer wavelength required for 3p excitation is significantly lower, resulting in a longer EAL^42–44^.

These advantages, however, come at the cost of a significantly higher required laser pulse energy, both due to the orders-of-magnitude lower 3p absorption cross-sections (over 37 orders, although units are different), and the higher water absorption at typical 3p excitation wavelengths for green indicators such as GCaMP^44^. As a result, the imaging depths obtained thus far (∼1.2 mm^37^) have been limited either by low fluorescence generated at the focus or laser-induced tissue heating, rather than the fundamental limit, set by a loss of SBR due to out-of-focus fluorescence. Moreover, efforts to maximize the imaging depth within the biological limits of the sample and the photophysics of 3p excitation have been hampered by ad hoc choices of the laser repetition rate, pulse energy, and pulse duration, combined with suboptimal pixel excitation schemes.

Here, we demonstrate Three-photon Ultra-Deep Imaging (TrUDI), a system which overcomes these limitations and reaches depths that approach the fundamental SBR depth limit of 3pM in the mouse brain. We have developed a 3p acquisition strategy in which excitation parameters of the laser and the scanning, as well as the optical parameters of the microscope, are optimally matched to linear and non-linear limits for safe laser exposure of the sample. Additionally, we have accounted for the spatial and temporal resolution requirements for single-cell resolution calcium (Ca^2+^) imaging, the photophysics of 3p excitation, and the scattering properties of mouse brain tissue. At comparable imaging depths, our approach yields a ∼250-fold increase in detected fluorescence photons at the same and typical laser power used in previous 3pM approaches.

These innovations allowed, for the first time, recordings of neuronal population activity at depths up to ∼2 mm in mouse brain tissue. We demonstrate this capability of our imaging system by performing imaging throughout the mouse motor cortex and into the striatum, as well as through the posterior parietal cortex (PPC) into the various regions of the mouse hippocampus, including the CA1 layer, the CA3 layer, and the dentate gyrus (DG) in awake mice. Throughout our recordings we maintained cellular resolution at frame rates of 5–10 Hz within a field of view (FOV) of ∼230 × 230 µm^2^ and at average power levels < 250 mW at the sample surface, which have previously been established to lie within safe limits for laser-induced tissue heating. This new parameter space and acquisition modality for 3pM open up new possibilities for capturing cellular-resolution dynamics of defined neuronal populations across a tissue column at previously unavailable depths within the scattering mammalian brain.

## Results

### Depth limit in three-photon microscopy

In multiphoton microcopy—when applied to the mammalian brain—the number of photons at the focus decreases exponentially with depth due to tissue scattering and absorption. To compensate for this loss of photons, and to maintain signal at the focus, the number of photons entering the sample needs to be increased exponentially with imaging depth which causes out-of-focus fluorescence at the superficial layers. Thus, the ultimate depth reach of multiphoton microscopy as characterized by the signal-to-background ratio (SBR) ^23,33–35^ is set by the depth limit at which the background fluorescence becomes equal to the fluorescence generated at the focus, rendering the SBR equal to unity.

**Fig. 1a** shows the dependence of SBR on focal depth *z_f_* in mouse brain tissue for 2pM at 920–960 nm, assuming an EAL 175 ± 25 μm^32,36,45–47^, and for 3pM at 1300 nm with an assumed EAL of 300 ± 50 µm^39,43,45^ (**Supplementary Note 1**), with the dashed line indicating the corresponding background-induced depth limits *z_DL_* (920 − 960 nm) = 0.6 ± 0.1 mm and *z_DL_* (1300 nm) = 2.3 ± 0.5 mm for when the 2p and 3p SBRs reach unity. Here we assumed that fluorescence is generated by ballistic photons only^23,35,36^, that the tissue is fluorescently labeled uniformly and that it has uniform scattering and absorption properties across the depth range, as characterized by a single EAL value. Accordingly, the indicated ranges for the EALs of the corresponding imaging modality reflect the observed variations from the above assumptions under realistic conditions. Thus, while the above result of *z_DL_* (920 − 960 nm) = 0.6 ± 0.1 mm is consistent with the generally reported experimental depth reach of 2pM, a major gap remains between the currently demonstrated depth reach of 3pM of ∼1.2 mm in Ca^2+^ imaging and the fundamental depth limit of ∼2.3 mm, as set by the level of out-of-focus fluorescence. As we discuss and demonstrate below, this gap can be accounted for by the lack of sufficient fluorescence signal at the focus as a result of sub-optimal excitation schemes and limitations set by laser-induced tissue heating.

**Fig. 1:**
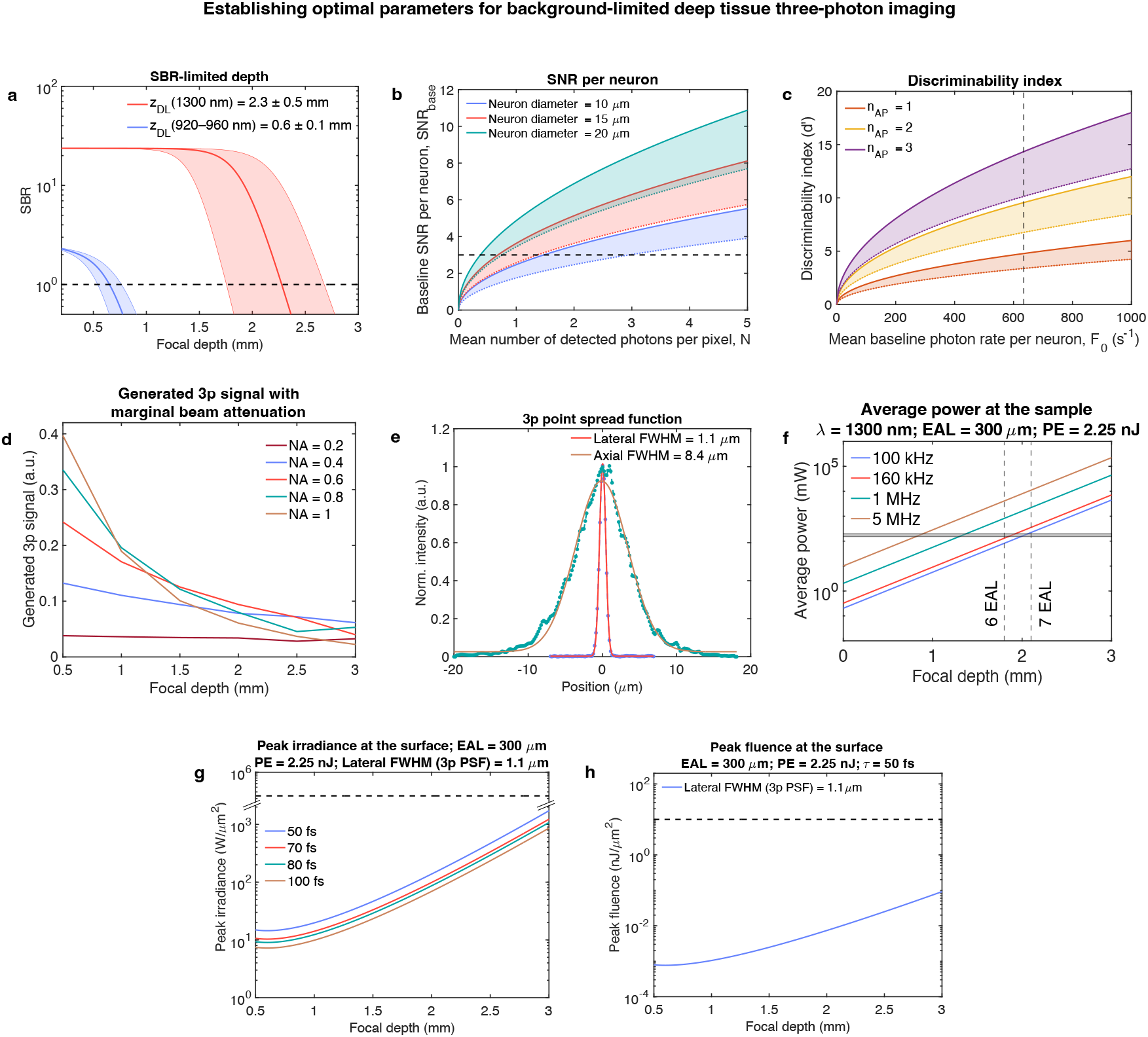
Establishing optimal parameters for background-limited deep-tissue three-photon imaging. (a) Signal-to-background ratio (SBR) as a function of imaging depth for two-photon fluorescence microscopy (2pM) at 920–960 nm (effective attenuation length, EAL 175 ± 25 µm) and three-photon fluorescence microscopy (3pM) at 1300 nm (EAL 300 ± 50 µm) in mouse brain tissue, assuming uniform labeling and attenuation properties with expected SBR-limited imaging depth ∼0.6 ± 0.1 mm for 2pM and 2.3 ± 0.5 mm for 3pM, where the SBR reaches unity. The shaded region represents the range of EAL variation. (b) Baseline SNR per neuron (SNR_base_) sampled by *M_pix_* pixels and hence excitation spots per neuron, as functions of the mean number of detected photons per pixel at the PMT, where fluorescence photons in each pixel are generated by a single pulse. The SNR plots are shown for three different neuron sizes (different color for 10, 15, and 20 µm diameter) for both conditions where background is negligible (i.e., SBR >> 1, upper solid lines of the shaded regions) as well as at SBR-limited depth (i.e., SBR = 1, lower dotted lines of the shaded regions). The horizontal black dashed line represents the minimum baseline SNR criterion of 3 per neuron. In **Extended Data** Fig. 1a**-b**, the required SNR_base_ threshold and the corresponding variance of the PMT output current per pixel are shown as functions of mean number of detected photons per pixel for the desired SNR_base_ = 3 per neuron. The theoretical model and the parameters are provided in **Supplementary Note 3**. (c) Discriminability index (*d*′), a measure of neuroactivity SNR, as a function of mean baseline photon rate per neuron (per second) for different numbers of action potentials (*n_AP_*), *n_AP_* = 1–3. The plots of *d*^′^ are shown for both conditions where background is negligible (i.e., SBR >> 1, upper solid lines of the shaded regions) as well as at SBR-limited depth (i.e., SBR = 1, lower dotted lines of the shaded regions). *d*^′^ provides a measure of the accuracy of neuronal activity event detection in terms of true positive rate (TPR) and false positive rate (FPR), which are also shown in **Extended Data** Fig. 1c**-d**, respectively, for SBR-limited depth. The vertical dashed line represents the mean baseline photon rate per neuron sampled with 25 pixels at a rate of 8.6 Hz in order to achieve a minimum baseline SNR threshold of 3 per neuron. (d) Generated 3p fluorescence signal as a function of focal depth for different excitation NAs. The fraction of ballistic photon reaching the focus and contributing to signal generation was modeled using Monte-Carlo simulation of photon propagation followed by nonlinear 3p fluorescence signal generation, assuming an EAL of 300 µm at 1300 nm and a paraxial objective lens with 12.5 mm focal length. For comparison, **Extended Data** Fig. 1e provides the generated 3p signal as a function of focal depth without this effect, for same excitation NA range. (e) Measured lateral (1.1 µm) and axial (8.4 µm) full-width-half-maximum (FWHM) of three-photon (3p) Point Spread Function (PSF) for the optimized deep tissue excitation using 0.5 µm green fluorescence beads at 1300 nm. (f) The required average power for 3pM as function of imaging depth for different laser repetition rates for a fixed pulse energy (PE) of 2.25 nJ at the focus, a typical value for generating sufficient fluorescence photons and assuming an EAL of 300 µm. The horizontal dashed line represents the limit for safe average power exposure of 250 mW^37,58^. The vertical dashed line denotes the focal depth at which a 160 kHz laser will reach the safe limit of average power to deliver 2.25 nJ pulse energy at the focus. (g) The peak irradiance at the surface as a function of focal depth for a PE of 2.25 nJ, a PSF size as shown in (e), and an assumed EAL of 300 µm at 1300 nm. Different colored plots correspond to four different pulse durations. The horizontal dashed line corresponds to the peak irradiance limit of ∼2.35 × 10^5^ W/µm^2^ for safe exposure^39^. (h) The peak fluence at the surface as a function of focal depth for the PSF size shown in (e), PE of 2.25 nJ, and pulse duration of 50 fs (FWHM), assuming an EAL of 300 µm at 1300 nm. The horizontal dashed line corresponds to the peak fluence limit of 10 nJ/µm^2^ for safe exposure^39^.

### Optimal excitation source and scanning parameters to reach the SBR-limited depth in of 3pM

Traditionally, 2pM but also a number of instantiations of 3pM have employed high-repetition-rate femtosecond lasers. Combined with the typical scan speed of galvanometric mirrors, this resulted in a condition where each image pixel is excited by multiple laser pulses. In this regime, the generated fluorescence signal scales linearly with the number of pulses per pixel. As we have previously shown^37,46,48,49^, using a laser at a lower repetition rate (*f_R_*_2_) to deliver the sum of the energy of *p* pulses of a laser with a higher repetition rate *f_R_*_1_ onto a single pixel leads to an effective increase in generated fluorescence signal by a factor 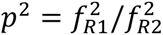 — in other words, a *p* times higher signal-to-noise ratio (SNR) at the same average power (**Supplementary Note 2**). Thus, a single-pulse-per-pixel excitation strategy allows for a more efficient use of the total available safe average power budget at the sample, thereby enabling an increase in FOV, acquisition frame rate, or in the case of 3pM, an increase of in-focus fluorescence signal at depth^37,46^. A few recent studies have utilized higher-pulse-energy, lower-repetition-rate lasers in the 0.4–1 MHz regime^32,36–39,50,51^, including some with the above single-pulse-per-pixel excitation scheme^37^. However, the exact choices of laser repetition rate and acquisition parameters appear to be somewhat ad-hoc, rather than optimized systematically for maximizing the obtainable imaging depth under safe biological imaging conditions. In the following, we discuss a principled approach that informed our bottom-up engineering and optimization scheme of all acquisition and laser parameters.

In the absence of noise, the SBR, and thus the obtainable depth limit, is independent of the pulse energy. However, in reality various types of noise (biological, readout, shot noise) require that the SNR for both the baseline fluorescence of neurons and their activity transients exceed a certain minimum, as discussed below. First, in order to detect the baseline fluorescence reliably, the noise of the measurement system (i.e., shot noise and systems noise; any noise associated with the signal detection process) needs to be overcome. To estimate the minimum number of photons per neuron per second required to achieve a baseline SNR (SNR_base_) of 3, we considered the typical range of mouse neuron soma diameters (∼10–20 μm^52–54^), as well as the need for their faithful spatial sampling and our inferred optimal choice of PSF (see below). Together with the single-pulse-per-pixel excitation principle, these dictate a minimum spatial sampling with ∼5 pixels in each dimension, or ∼25 pixels in total, per neuron. We assumed Poisson-distributed photons at the detector at a constant mean rate and included an additional noise component that is contributed by the photomultiplier tube (PMT) gain in the regime of low average photon numbers. This allowed us to establish the relationship between the anode current, its variance (or equivalently, pixel digital count and count variance) and the mean number of detected photons (**Supplementary Note 3**). For typical parameters of our system (PMT gain: 2 × 10^6^, quantum efficiency: 0.45, cathode radiant sensitivity: 189 mA/W, fluorescence response width: 3.5 ns), the SNR_base_ of a neuron is shown in **Fig. 1b** as functions of mean number of detected photons per pixel, for different neuron sizes for both, in the absence of fluorescence background as well as at the SBR limit of 1. In **Extended Data Fig. 1a-b**, the required SNR_base_ threshold and the corresponding variance of the PMT output current per pixel are shown as functions of mean number of detected photons per pixel for the desired SNR_base_ = 3 per neuron. For our smallest (largest) assumed neuron size of ∼10 μm (20 μm) diameter, sampled by 25 (100) pixels, the required baseline pixel SNR (SNR_base,pix_) is ∼0.6 (0.3) per frame, respectively. At the SBR-limited imaging depth, this corresponds to ∼3 (0.8) detected photons per pulse or – irrespective of the neuron size – to ∼635 detected photons per neuron per second at our chosen framerate of 8.6 Hz (see below and **Supplementary Note 3**) while for the depth range (≲1 mm) where the fluorescence background is negligible this requirement drops to ∼320 photons per neuron per second.

The discrimination of neurons’ Ca^2+^ activity from their baseline fluorescence can be characterized by the discriminability index (*d*′) given by 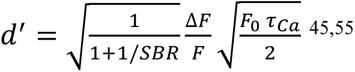, where 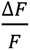 is the relative fluorescence change induced by one action potential (AP), *F*_0_ the baseline fluorescence photon detection rate and *τ_Ca_* the half-life of the Ca^2+^ decay (**Supplementary Note 4**). If multiple APs occur within *τ_Ca_*, the observed 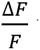 will scale approximately linearly with the number of APs (*n_AP_*) for small (∼1–3) *n_AP_*^30^. Thus, the AP-dependent discriminability index can be written as 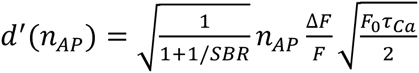. We note that *d*^′^ is proportional to the z-score (obtained by subtracting the baseline mean from the raw fluorescence time trace and dividing the result by the baseline standard deviation), another measure of neuronal activity. Both *d*^′^ and the z-score depend on the baseline SNR_base_ via the factor = 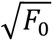. We evaluated *d*^′^(*n_AP_*) under the conditions that we had previously determined to yield a baseline SNR_base_ greater than 3. At the background-limited depth reach, corresponding to an SBR of 1, using the *in vivo* parameters of GCaMP8s^30^ (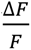 = 0.6 and *τ_Ca_* = 0.2 *s*) and at our imaging frame rate, we found *d*^′^(*n_AP_*) to be 3.4, 6.8 and 10.1 for 1, 2, and 3 APs respectively. This corresponds to a Ca^2+^ transient detection rate (true positive rate) between > 95.5–99.9%, and a false positive rate of < 4.5–0.1% for 1 to 3 APs, respectively. In **Fig. 1c**, the discriminability index is plotted for 1–3 APs as a function of the detected mean baseline photon rate per neuron, and in **Extended Data Fig. 1c-d**, the corresponding true positive rate (TPR) and false positive rates (FPR) are shown (**Supplementary Note 4**). Thus, we found that the requirement of SNR_base_ = 3 was sufficient to ensure that the requirement of *d*^′^ = 3 for detecting Ca^2+^ transient events at the SBR limited depth is fulfilled.

Next, we aimed to identify the optimal size of the excitation point spread function (PSF) that maximizes the number of generated photons for a given pulse energy. Traditionally, using the highest possible numerical aperture (NA) for both excitation and photon collection has been assumed to maximize signal under non-saturation conditions. However, when imaging neuronal cell bodies at depth in scattering tissue, we found that this assumption requires reevaluation with respect to the excitation NA, for two reasons: First, the signal generated by the excitation PSF is proportional to the cube of the photon flux and linearly proportional to the axial extent of the PSF in 3pM. Thus, a larger axial PSF, as long as it does not exceed the average diameter of neuronal cell bodies, could enable integration of the fluorescence signal over a greater number of fluorophores. It has been shown that for neuronal imaging, an axial extent comparable to the size of the cell body diameter (∼10–15 µm) is preferable^35,37^. Second, a higher excitation NA means that the peripheral rays have a longer beam path to the focal point. This translates to a greater degree of scatter-induced loss of excitation photons at the focus and therefore a reduction of fluorescence signal generation in 3p with increasing imaging depth compared to a lower excitation NA. **Extended Data Fig. 1e** illustrates the generated 3p signal as a function of focal depth without this effect for excitation NAs of 0.2–1.0, showing the expected increase in 3p signal as function excitation NA for constant pulse energy at the focus, independent of imaging depth. We then took the effect of scatter-induced loss of photons for marginal rays into account by performing Monte-Carlo simulations to estimate the fraction of ballistic photons that reached the focus and contributed to the generated 3p fluorescence signal, as a function of focal depth and for different excitation NAs, assuming the same pulse energy at the tissue surface (**Fig. 1d**). We found that at lower tissue depth (≲1 mm), the traditional expectation that the generated fluorescence signal is maximized for the highest excitation NAs holds. However, at greater tissue depths the optimal excitation NA was lower, as a result of a higher probability of loss of ballistic photons in the marginal rays due to an effectively longer travel path of excitation photons, and hence higher non-linear falloff of the generated fluorescence signal, at larger NAs. Specifically, we found that the optimal excitation NA at the SBR-limited depth of ∼2–3 mm in the mouse brain is ∼0.4–0.6. We note that while this lower NA range leads to optimal excitation, it is still beneficial to use the full available NA of a high-NA objective for collection (NA 0.8 in our case).

This finding is consistent with other reports; for example, Wang *et al*. suggested that for imaging depth of 6–7 EAL in 3p, the optimal fill factor for high NA objective drops to about 50%^56^ (which is equivalent to an NA ∼ 0.4 for an objective of NA of 0.8). In practice, the signal loss at high NAs can be even more severe than what is accounted for by the longer path length of marginal rays. For instance, Dunn *et al*. showed that the optimal NA for 2p excitation in scattering tissue decreases from 1.4 to ∼0.6 as imaging depth increases from 100 μm to 300 μm^57^. In addition, we note that the beam diameter at the surface scales as ∼NA × 3 for ∼2 mm imaging depth (beam diameter ∼ 2 *z_f_* sin *θ*, where sin *θ* = NA/*n_idx_* ; *z_f_* = 2 mm, the imaging focal depth; *n_idx_* = 1.33, the refractive index). Thus, signal attenuation is indirectly affected more strongly at higher NAs by factors such as surface inhomogeneities, larger blood vessels, and aberrations. Therefore, a lower effective excitation NA is advantageous for reaching the SBR-limited depth of 6–7 EALs in 3p. Based on these considerations, we used a lateral 3p PSF size of ∼ 1.1 µm full-with-at-half-maximum (FWHM) and an axial 3p PSF size of 8.4 µm FWHM in our TrUDI design (**Fig. 1e**).

As the next step, we sought to identify the required pulse energy based on this optimal PSF size and the number of photons needed per neuron per second to achieve a desired *d*^′^ for reliable Ca^2+^ transient detection. As shown above, a minimum of 635 detected photons per neuron per second is required to satisfy both SNR_base_ = 3 as well as a minimum *d*^′^ of 3 for detecting Ca^2+^ transient events. This corresponds to ∼3.0 detected photons per pulse, assuming sampling of the smallest neurons cell bodies (diameters ∼10 µm) with 25 pulses at a frame rate of 8.6 Hz. Using our experimentally informed model for fluorescence generation and collection as well as the system noise (**Supplementary Note 3**) we found that to achieve this, a pulse energy of ∼2.25 nJ at the focus is sufficient at our PSF size. We then corroborated our findings regarding the optimal PSF size and required pulse energy by performing *in vivo* experiments and evaluating the experimentally obtainable pixel SNR. We found good agreement with our theoretical model for SNR (**Supplementary Note 3**), both for the experimental variance of pixel photon count (**Extended Data Fig. 2a**) and the pixel SNR (**Extended Data Fig. 2b)** as a function of mean pixel count. Furthermore, the histogram of our experimentally obtained pixel SNR (**Extended Data Fig. 2c**) for ∼2 nJ pulse energy at the focus demonstrates that our criterion for the baseline SNR per pixel (**Extended Data Fig. 1a**) was satisfied. These findings are consistent with previous reports of 3pM in which pulse energies of ∼1–5 nJ at the focus (for ∼40 fs pulses) resulted in sufficient fluorescence signal and SNR for reliable Ca^2+^ imaging while remaining well below the ∼10 nJ/µm^2^ peak fluence threshold for non-linear damage^39^.

Next, we investigated the relationship between laser repetition rate, average power and obtainable depth for a pulse energy of 2.25 nJ at the focus. **Fig. 1f** shows the relationship between the required average power and the obtainable imaging depth when delivering 2.25 nJ energy at the focus while staying within the safe average power limit of ∼250 mW^58^ (assuming EAL 300 µm and 50 fs pulse duration, for different laser repetition rates). This plot illustrates the key roadblock in reaching the fundamental SBR-limited 3pM imaging depth when MHz range repetition rate lasers are used: With these repetition rates, it is impossible to reach the SBR-limited depth because delivering the required pulse energy at the focus would make it necessary to exceed the average power safety limit by more than an order of magnitude. For example, in a previous study that implemented one pulse per pixel excitation, in order to deliver ∼3.5 nJ at the focus ∼190 mW of power was required for an imaging depth of ∼1.2 mm using a ∼1 MHz, 70 fs laser^37^. At this repetition rate, to reach a depth of 2 mm while keeping the pulse energy at the focus constant, >2.5 W of average power would be needed. Clearly, to approach the fundamental SBR-limited imaging depth of 3pM at safe average power while delivering sufficient pulse energy at the focus, lower laser repetition rates are required. As shown in **Fig. 1f**, a 160 kHz laser can deliver 2.25 nJ pulse at ∼ 2 mm depth while keeping the average power at the surface below ∼250 mW. Since the generated fluorescence signal in 3pM is proportional to the cube of the pulse energy, this corresponds to a ∼250-fold increase in 3p excitation efficiency at a given average laser power.

However, a few additional considerations are required when using such low repetition rate lasers: First, as discussed above, in order to spatially resolve neurons and their boundaries, each neuron needs to be sampled by ∼25 separated spots^37,46,49^. Thus, the pixel spacing needs to be selected appropriately to ensure faithful spatial sampling of each neuron while maximizing the imaging FOV. For the smallest neurons we considered, with a soma diameter of 10 µm, this translated to pixel spacing of ∼1.8 µm. Second, reducing the laser repetition rate, while keeping all other parameters constant, inevitably decreases the pixel acquisition rate and thus the frame rate. This necessitates balancing the tradeoff between spatial sampling and FOV with the additional requirement of sufficient temporal resolution for *in vivo* Ca^2+^ imaging. Third, as the pulse energy increases, the peak fluence and—depending on the pulse duration—the peak irradiance can in principle exceed the safety limits for non-linear laser exposure at the tissue surface. Assuming again a pulse energy of ∼2.25 nJ at the focus, **Fig. 1g-h** show the resulting peak irradiance and the peak fluence at the sample surface as a function of focal depth. The safe limits for fluence and irradiance of ∼10 nJ/µm^2^ and ∼2.3×10^5^ W/µm^2^, respectively^39^, would be exceeded at a hypothetical imaging depth of ∼5 mm (**Extended Data Fig. 3a-b**). However, for the ultimate obtainable imaging depth of ∼2.3 mm in 3pM, as set by the SBR > 1 requirement, both the fluence and the irradiance remain well within safe limits at the surface, and much more so at the focus (**Fig. 1a**). Finally, as shown in **Fig. 1g**, since our identified optimal excitation parameter space allows to remain well within the safe tissue irradiance thresholds for pulse durations over the entire a range of 50–100 fs, and since the excitation efficiency in 3pM scales as the inverse squared of the pulse duration^37,39,45^ (**Supplementary Note 1**), we maximized the obtainable fluorescence signal for the same average power by choosing the shortest possible pulse duration of ∼ 50 fs while maximizing the overlap of the resulting spectral full bandwidth of ∼85 nm with the 3p absorption band near the absorption peak of GCaMP^59^.

To experimentally implement the inferred laser pulse parameters and optimal excitation conditions for deep 3pM, we used a customized tunable optical parametric amplifier (OPA) outputting ∼50 fs pulse with > 10 µJ at 160 kHz (**Methods**). Mapping this repetition rate onto the above requirements for spatial sampling of neurons and the temporal dynamics of GCaMPs informed the design of our TrUDI system. TrUDI provides access to a FOV of ∼230 × 230 μm² at a pixel spacing of 1.8 μm while operating under a single pulse per pixel excitation scheme and at a frame rate of 8.6 Hz (**Methods**). Thus, it allows for imaging of neuronal dynamics at unprecedented tissue depths in the mouse brain, at cellular and subcellular-scale spatial resolution, within a moderate FOV and at adequate temporal resolution. A schematic of the optical path of the TrUDI is shown in **Extended Data Fig. 4**.

### Experimental comparison of TrUDI with MHz 3pM and in vivo validation of imaging performance

To experimentally validate our optimization ideas, we performed a direct comparison of TrUDI with a state-of-the-art 3pM implementation that used one-pulse-per-pixel acquisition at 1 MHz repetition rate, ∼70 fs pulse duration and ∼3.5 nJ pulse energy. This reference implementation had previously achieved the deepest 3p Ca^2+^ imaging to date, reaching ∼1.2 mm depth at a mean SNR_base_ of ∼2.5^37^. We matched PSF and detection parameters to our TrUDI setup and constructed a similar optical path, with the exception of a longer pulse duration of ∼100 fs, necessitated by experimental constraints (**Methods**). We then performed a comparative *in vivo* imaging in the PPC in mice expressing GCaMP6s (Camk2a-tTA × tetO-GCaMP6s) and sought to reach the same imaging depth in the same animal both using the 1 MHz reference system and TrUDI (**Fig. 2a-b**). While the 1 MHz system required 193 mW of average power at the sample surface to achieve a mean SNR_base_ of 2.2 at a depth of 820 µm, TrUDI reached the same imaging depth with only 7.5 mW of average power while achieving a mean SNR_base_ of ∼6.1. This is consistent with our numerical estimates that TrUDI’s lower repetition rate leads to a ∼250-fold increase in 3p excitation efficiency.

**Fig. 2:**
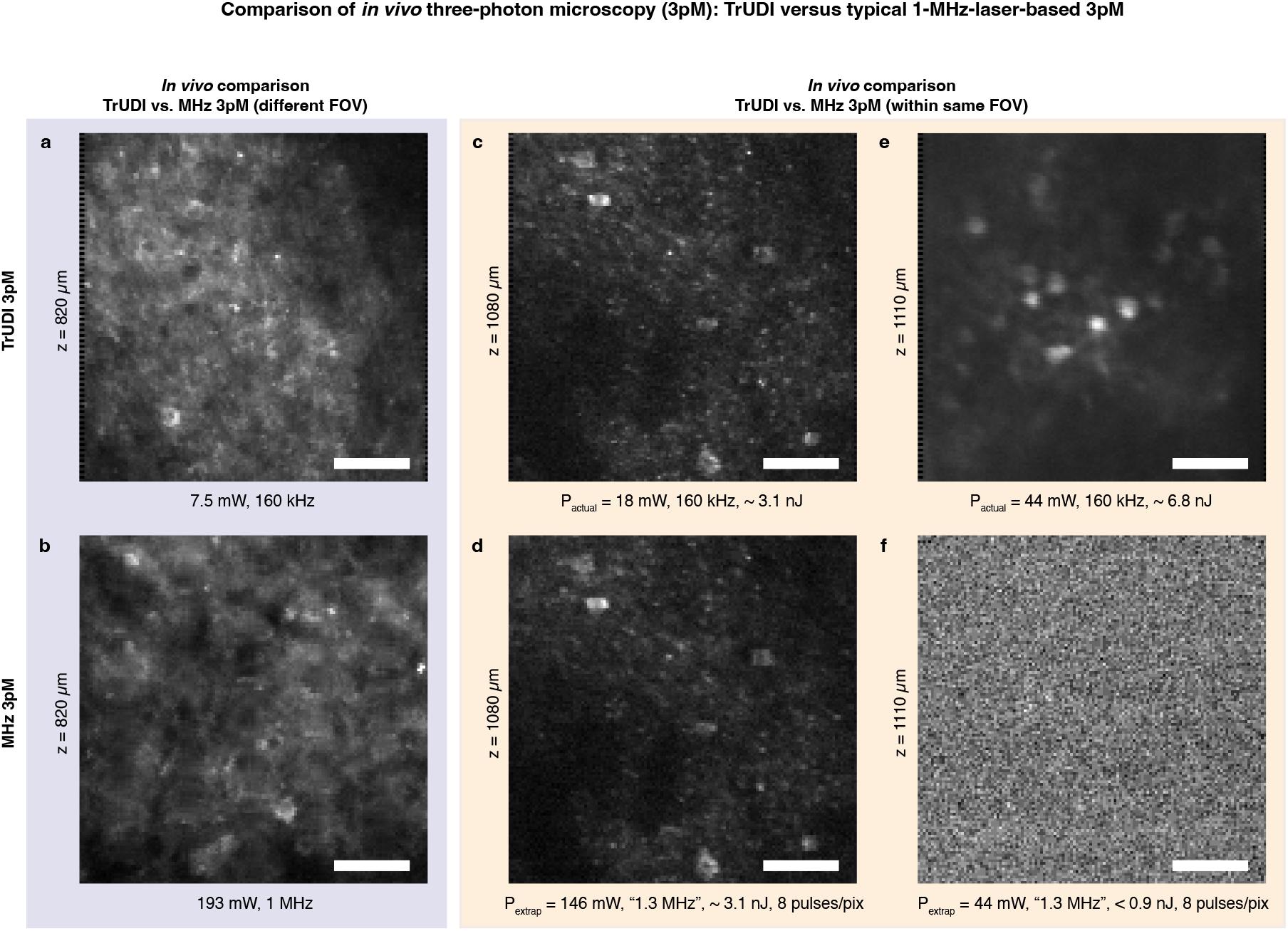
Comparison of *in vivo* three-photon microscopy (3pM): TrUDI versus typical 1-MHz-laser-based 3pM. (**a-b**) Comparison of *in vivo* imaging between TrUDI and a 1-MHz-laser-based 3pM to reach an imaging depth of 820 µm within the posterior parietal cortex (PPC) in mouse expressing GCaMP6s: (a) TrUDI’s 160 kHz system required an average power of ∼7.5 mW (∼3 nJ at the focus), (b) 1 MHz system required 193 mW (Note: TrUDI had a pulse duration of ∼50 fs while that of the 1 MHz system was ∼100 fs.). Scale bar, 50 µm. (**c-d**) Comparison of the required average power between TrUDI and 1-MHz 3pM configuration for a fixed pulse energy within the same imaging FOV for similar pixel SNR at 1080 µm depth in hippocampus (HPC) of a mouse expressing GCaMP8m. (c) TrUDI required an average power (*P_actual_*) of 18 mW (∼3.1 nJ at the focus), (d) the 1 MHz configuration required an average power (*P_extrap_*) of ∼146 mW for the same pulse energy at the focus. (see **Methods** and main text). Scale bar, 50 µm. (**e-f**) Comparison of the SNR between TrUDI and 1-MHz 3pM configuration using the same average power within the same imaging FOV at 1110 µm depth in HPC of a mouse expressing GCaMP8m. (e) TrUDI with its 160 kHz repetition rate yields clearly resolvable neurons with a mean baseline SNR (SNR_base_) of ∼ 4.8 using an average power (*P_actual_*) of ∼ 44 mW while the SNR_base_ for the 1 MHz configuration using the same extrapolated average power, (*P_extrap_* = 44 mW) is too low to resolve neurons, as shown in (f). Scale bar, 50 µm.

In addition, to eliminate potentially confounding differences between microscopes and enable a direct comparison within the same apparatus, animal, and 3D FOV, we mimicked the reference 1 MHz imaging conditions on the TrUDI optical path. Since operating TrUDI’s laser system in the MHz regime was not possible, we used its 160 kHz laser to spatially oversample the FOV by slowing down the scanner, effectively emulating a 1.3 MHz system. The use of the same pulse energy in both imaging modalities led to an 8-fold higher “extrapolated” average power (*P_extrap_*) for the 1.3 MHz configuration compared to the actual average power (*P_actual_*) used in TrUDI. Moreover, to keep the pixel pitch (1.8 µm) constant between the two modalities, we averaged the signal from 8 successive pulses into a single pixel. Using this approach to mimic the reference approach, we compared it to TrUDI under two different conditions: one in which the pulse energy was kept the same (**Fig. 2c** vs. **Fig. 2d**), and another in which the average power (*P_actual_* and *P_extrap_* for TrUDI and the 1.3 MHz configuration, respectively) was kept the same in both systems, (**Fig. 2e** vs. **Fig. 2f**).

In the first scenario, we performed imaging in the hippocampus through intact cortex at a depth of 1080 µm using TrUDI, and found that only ∼18 mW of average power (*P_actual_*) at the sample surface was sufficient to resolve neurons with estimated mean SNR_base_ of ∼2.2 (**Fig. 2c**). This corresponds to a pulse energy of ∼ 3.1 nJ at the focus, similar to the energy used in other 3p imaging studies^37,39,45,60^. In contrast, imaging at the same depth with the same pulse energy in the 1.3 MHz configuration required ∼146 mW of average power (*P_extrap_*) (**Fig. 2d**). Thus, MHz-rate 3pM systems would well exceed the safe exposure limit at the SBR-limited imaging depth (> 2 mm).

In the second scenario, we used ∼ 44 mW of average power (*P_actual_*) at the surface in TrUDI, which resulted in clearly resolved individual neurons at 1.1 mm depth (mean SNR_base_ ∼ 4.8) (**Fig. 2e**). To maintain the same average power in the 1.3 MHz configuration (*P_extrap_* = 44 mW), the pulse energy needed to be reduced 8-fold to <0.9 nJ which resulted in a ∼500-fold lower 3p signal and thus prohibitively low SNR_base_ to identify and segment neurons (**Fig. 2f)**.

These results demonstrate the potential of TrUDI for extending the obtainable imaging depths in the scattering mouse brain while requiring minimal average power. This is achieved by combining a lower-repetition-rate laser with a 3p acquisition system optimized to maximize in-focus fluorescence signal generation per pulse. We next explored the maximum *in vivo* imaging depth achievable with TrUDI and its ability to record neuronal population activity at unprecedented tissue depths in the mouse brain.

### Structural imaging and neuronal population recording in the mouse hippocampus through intact cortex at 1730 µm depth

Our initial *in vivo* characterization already demonstrates the practical gains in in-focus fluorescence signal achievable with TrUDI. Thus, we expected that it would enable both, higher-SNR imaging at the previously demonstrated tissue depths at lower average power, and imaging at previously unattainable tissue depths within the safe average power limits. To demonstrate this, we imaged through the cortical layers in the posterior parietal cortex (PPC) of mice expressing green fluorescent protein (Thy1-EGFP) into the hippocampus (HPC) region (**Fig. 3a-b**). A high-resolution reconstruction of the acquisition volume using an axial plane spacing of 5 µm and extending to 1200 µm depth is shown in **Fig. 3b**, with selected single planes shown in **Fig. 3c-h**. Detailed neuronal processes were observed throughout the PPC and the dorsal HPC, using ∼76 mW at the deepest imaging plane at ∼1200 µm, beyond the corpus callosum (CC). Additionally, we found highly detailed neurite structures associated with the CA1 pyramidal layer within a depth range of 1050–1250 µm (**Fig. 3i-m**). Our key motivation, however, for optimizing the in-focus fluorescence signal was to increase the obtainable imaging depth towards the SBR-limited range (∼ 2.3 mm). Under these conditions, individual neurons were indeed clearly visible with a high SNR_base_ (> 7.0) at depths up to 1.73 mm, using ∼105 mW of average power, well below established thresholds for thermal damage^37,39,45,58^ in multiphoton microscopy (**Fig. 3n**).

**Fig. 3:**
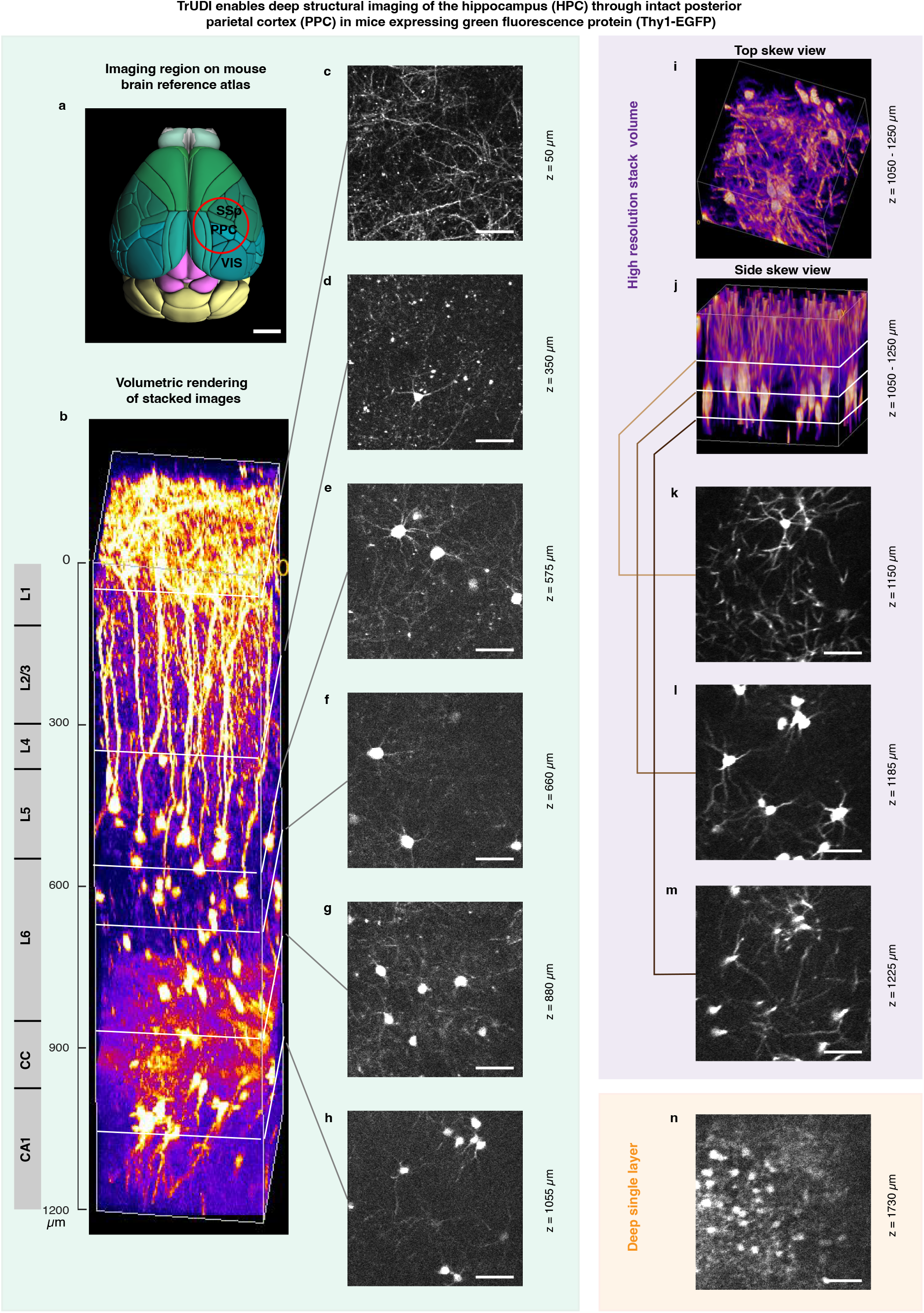
TrUDI enables deep structural imaging of the hippocampus (HPC) through intact posterior parietal cortex (PPC) in mice expressing green fluorescence protein (Thy1-EGFP). (**a**) Rendering of the mouse brain based on the Allen mouse brain atlas showing the approximate location of a 4-mm cranial window (red circle) over the PPC and the subcortical HPC, from which the recordings in panels (b-n) were obtained. Scale bar, 2 mm. (**b**) High-resolution static volume stack covering ∼1200 µm depth range showing detailed neuronal processes in the PPC and HPC through the corpus callosum (CC), acquired using between 0.7 and 76 mW of average power, with 76 mW being used for the deepest layer of CA1 in HPC. Mean intensity projection over 20 frames per plane were used. Size of the imaged volume: 230 × 230 × 1200 µm^3^ using 0.9 × 0.9 × 5 µm^3^ pixel spacing. (**c-h**) Single-plane slices of the volume in (b) are shown for selected depths at 50, 350, 575, 660, 880, and 1055 µm, with corresponding average acquisition power values of 0.8, 1.3, 2.0, 3.0, 14.0, and 54.0 mW, respectively. Scale bar, 50 µm. (**i-j**) Two 3D rendering views of high-resolution recordings of the dorsal hippocampus covering a depth range of 1050–1250 µm generated using an average power of 50 mW. Size of imaged volume: 230 × 230 × 200 µm^3^ using 0.9 × 0.9 × 5 µm^3^ pixel spacing. Scale bar, 50 µm. (**k-m**) Single-plane slices within the deeper range (∼1150–1225 µm) of volume shown in (j), revealing detailed views of dendrites, soma, and stems of axons. Scale bar, 50 µm. (**n**) An example single-plane recording at 1730 µm, a depth about ∼2 EAL greater than previously achievable limits^37,38^, acquired using 105 mW of average power, which is substantially lower than the established upper bound for safe power limits. Mean intensity projection over 20 frames. Size of imaged area: 230 × 230 µm^2^ using 1.8 × 1.8 μm pixel spacing. Scale bar, 50 µm.

### Neuronal population imaging in posterior parietal, primary visual cortex and hippocampus at up to 1840 µm depth

To validate the capability of our method to capture neuronal population activity at large tissue depth we imaged transgenic mice expressing GCaMP8s (Camk2a-tTA × tetO-GCaMP8s) or GCaMP8m (VGAT-Cre × GCaMP8m) throughout the cortex and sub-cortical regions (**Methods**). Using TrUDI, we recorded neuroactivity through various layers of the PPC or the primary visual cortex (V1), and further into the hippocampus, in awake, head-fixed mice during spontaneous activity (**Fig. 4a-b**). Using similar recording conditions and parameters as above, we captured the activity of neurons within a FOV of 230 × 230 μm² at a pixel spacing of 1.8 µm at 8.6 Hz frame rate. **Fig. 4b** and **Supplementary Video 1** show a 3D representation of our time-projected neuronal recordings across 170 planes, spaced axially spaced by 10 μm, covering the PPC and the HPC and reaching a maximal depth of ∼1.7 mm in a mouse expressing GCaMP8m (**Methods**). Consistent with our initial *in vivo* characterization results (**Fig. 3**), we found the maximum necessary average power for our deepest recordings to be ∼127 mW, well below the established limits for safe *in vivo* power exposure. We note that a significant part of the power increase occurred while passing through the CC, due to its strong scattering properties as a result of concentration of myelinated axons in this region. For comparison, the required power for imaging at 750 µm was only ∼ 4.7 mW, while imaging at 1030 µm required an average power of ∼21 mW. We performed 6–10-minute recordings of neuronal activity at 8.6 Hz across multiple planes ranging from 730 to1840 μm (**Fig. 4c-g**). The shown planes include deeper cortical layers in PPC as well as hippocampal CA1 and DG neuronal populations. We segmented neurons and extracted the time traces of their activity in each plane after performing motion correction (**Extended Data Fig. 5, Methods**), followed by processing of the data using our custom, CaImAn-based^61^ data reconstruction pipeline, which allowed for reliable segmentation and extraction of the neuronal footprints and their corresponding activity traces using a combination of spatiotemporal filters and non-negative matrix factorization (NMF) (**Methods**).

**Fig. 4:**
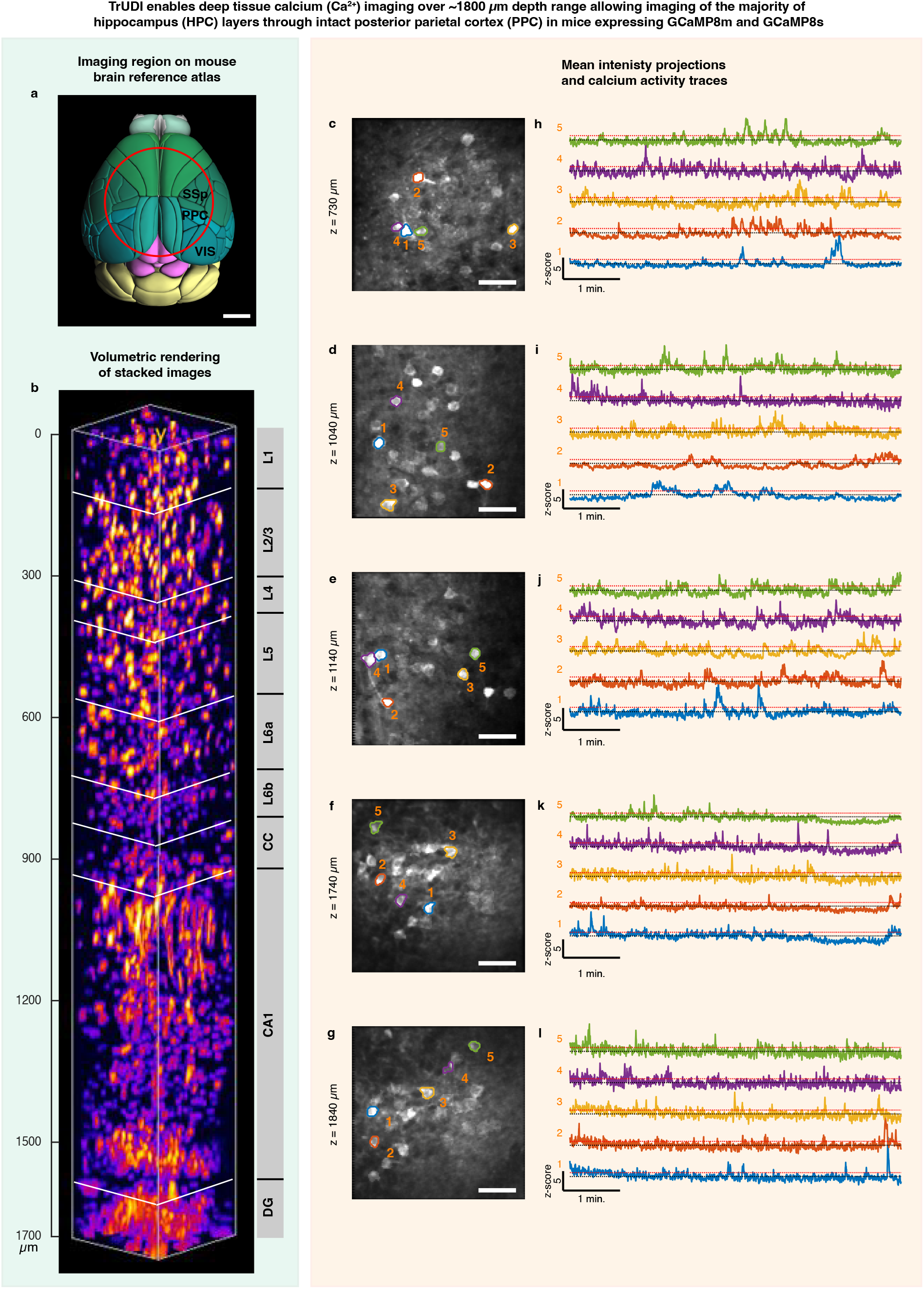
TrUDI enables deep tissue calcium (Ca^2+^) imaging over a ∼1800 µm depth range, allowing imaging of the majority of hippocampus (HPC) layers through intact posterior parietal cortex (PPC) in mice expressing GCaMP8m and GCaMP8s. (a) Rendering of the mouse brain based on the Allen mouse brain atlas showing the approximate location of an 8-mm cranial window (red circle) over the PPC and the subcortical HPC, from which the recordings in (b-l) were obtained. Scale bar, 2 mm. (b) High-resolution reconstructed volume stack covering a ∼1700 µm depth range, showing varying density of neurons across different layers (L1–L6) of the PPC, as well as the neurons in the hippocampal CA1 and DG region. The acquisition power was increased exponentially between from 0.4 to 127 mW to obtain constant signal across depths. Mean intensity projection based on recording of 1 minute duration at 8.6 Hz for each layer. Size of the imaged volume: 230 × 230 × 1700 µm^3^ using 1.8 × 1.8 × 10 µm pixel spacing. (**c-g**) Mean intensity projection of neuroactivity in individual planes covering a depth range of 730–1840 µm, showing clearly resolvable neurons and corresponding Ca^2+^ activity traces (h-l), even in the deepest recordings at 1840 µm. The average power used for each of the five depths was 2.5, 6.5, 12, 58, and 66 mW, respectively, all of which were significantly lower than the established upper bound for safe power limits. Imaging area, 230 × 230 µm^2^; pixel size of 1.8 × 1.8 µm^2^. Scale bar, 50 µm. (**h-l**) Ca^2+^ activity traces of selected neurons numbered in (c-g) over ∼6 min. using CaImAn^61^ with z-scores up to ∼ 0 for the strongest responding neurons. The red dotted line denotes a z-score of 1 and the black dotted line denotes the baseline.

Within each plane, neurons could be clearly identified and segmented, and their Ca^2+^ dynamics reliably extracted across the full depth range. **Fig. 4h-l** show example neurons and time traces from recordings within a depth range of 1040 to 1840 µm. **Extended Data Fig. 6** shows additional segmented neurons and their corresponding time traces in each plane, with **Supplementary Video 2** showing corresponding videos of single-plane neuroactivity at 1040, 1140, 1740 and 1840 µm, respectively. The maximum average power in these recordings was ∼66 mW at the deepest depth of 1840 µm, which was lower than what was needed for our Thy1-GFP mouse imaging (**Fig. 3n**) and the GCaMP8m (**Fig. 4b**) imaging at comparable depths. We believe this may be due to the higher brightness and sensitivity of the tetO-GCaMP8s indicator under the Camk2a-tTA promoter.

### TrUDI enables neuronal population imaging through motor cortex and the striatum at up to 1930 µm

Next, we validated the performance of TrUDI by recording neuronal population activity across depths and layers of the secondary motor cortex (M2) as well as the striatum (**Fig. 5a**) in head-fixed mice expressing GCaMP6s (Camk2a-tTA × tetO-GCaMP6s) during spontaneous activity. Using a 20 µm axial plane spacing and otherwise the same imaging parameters as in **Fig. 4b**, we recorded the activity of the neurons from the superficial layers of the motor cortex down to ∼2.1 mm depth from the pia. **Fig. 5b** shows the reconstructed volume of time-projected neuronal population activity, where active neurons are clearly visible across different depths. In this case, the maximally required average power was ∼150 mW for our deepest recordings at ∼2.1 mm. **Fig. 5c-g** show time projected images from 4–10-minute-long recordings of individual planes at 8.6 Hz, within a range of 1.05–1.93 mm. The data was processed through our motion correction and neuron segmentation pipeline similar to the data in **Fig. 4**. The time series corresponding to the indicated neurons are shown in **Fig. 5h-l**, displaying clear Ca^2+^ transients for example neurons through depths up to 1930 µm. **Extended Data Fig. 7** shows additional segmented neurons and corresponding time traces in each plane, with **Supplementary Video 3** showing corresponding videos of single-plane neuroactivity at 1050, 1400, 1810 and 1930 µm, respectively. The maximum required average power for these recordings was ∼120 mW for our deepest plane at 1930 µm.

**Fig. 5:**
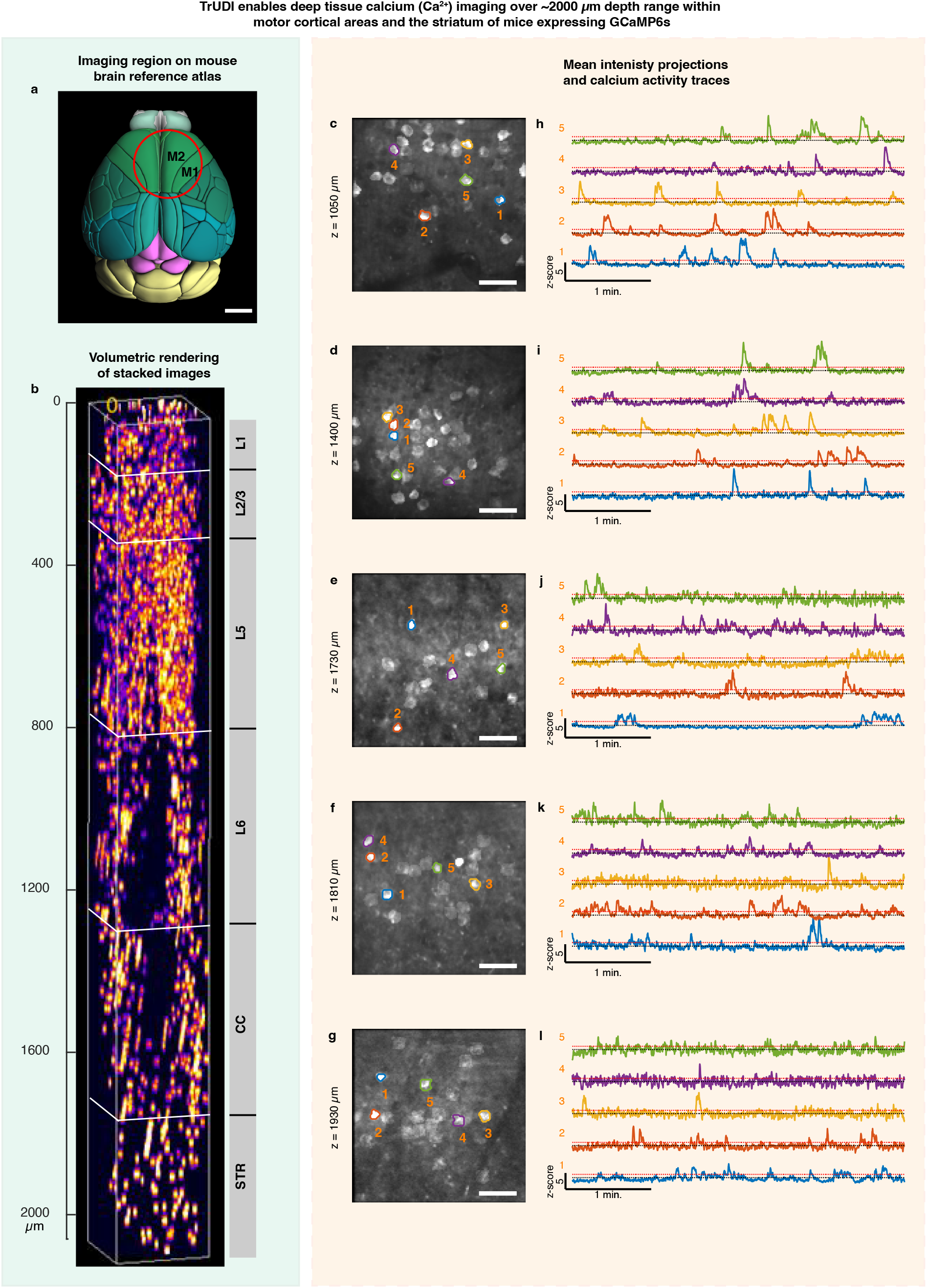
TrUDI enables deep-tissue calcium (Ca^2+^) imaging over ∼2000 µm depth range within motor cortical areas and the striatum of mice expressing GCaMP6s. **a**) Approximate location of a 5-mm cranial window (red circle) over the mouse brain, targeting deep imaging in the secondary motor cortical area (M2) and striatum, shown over the Allen mouse brain atlas. Scale bar, 2 mm. (**b**) High-resolution reconstructed volume stack covering a ∼2060 µm depth range showing varying density of neurons across different layers (L1–L6) of M2 as well as the neurons in the striatum. The acquisition power was exponentially increased from 0.5 to 150 mW to obtain constant signal across depths. Size of the imaged volume: 230 × 230 × 2060 µm^3^ using 1.8 × 1.8 × 20 µm pixel spacing. Images show mean intensity projections over ∼1 min at 8.6 Hz. (**c-g**) Mean intensity projection of neuroactivity in individual planes covering a depth range of 1050–1930 µm, showing clearly resolvable neurons and corresponding Ca^2+^ activity traces (h-l), even in the deepest recordings at 1930 µm. The average power used for each of the five depths was 10, 30, 120, 120, and 120 mW, respectively, all substantially lower than the established upper bound for safe power limits. Imaging FOV, 230 × 230 µm^2^; pixel size of 1.8 × 1.8 µm^2^. Scale bar, 50 µm. (**h-l**) Ca^2+^ activity traces of selected neurons numbered in (c-g) over ∼ 4 min. Traces were extracted from recordings using CaImAn^61^ and show z-scores up to ∼12 for the strongest responding neurons. The red dotted line denotes a z-score of 1 and the black dotted line denotes the baseline.

Our results clearly demonstrate the capabilities of our new imaging system. By performing systematic and careful optimization of excitation laser parameters, acquisition modality and the optical parameters of our 3pM system, we demonstrated a ∼250-fold effective increase in in-focus fluorescence signal within the established safe limits of average power and energy density. This enabled Ca^2+^ imaging across ∼6 EAL. To our knowledge, this represents the first demonstration of Ca^2+^ imaging over such a depth range. Our demonstrated imaging depth of ∼2 mm through intact brain tissue exceeds previous 3pM achievements by ∼800 µm—corresponding to ∼2–3 EALs— and approaches the fundamental SBR-limited imaging depth in 3pM at 1300 nm in the mouse brain.

## Discussion

In the last decade, 3pM has emerged as a powerful microscopy technique for neuroscience by extending the imaging depth reach beyond the ∼600 µm possible with 2pM in the scattering mammalian brain. While 3pM has found increased application in various areas of neuroscience, its key limitation is the unproportionally high heat penalty due to the higher-order nonlinearity compared to 2pM. Thus, all demonstrations of 3pM to date have been constrained by the in-focus signal achievable within safe average power limits. In contrast, our results show that without this limitation, the true SBR-ratio limited depth of 3pM at 1300 nm—the optimal wavelength for exciting green Ca^2+^ indicators such as GCaMP, is ∼2.3 mm.

In this work we show how the bottom-up principled optimization of all optical parameters of a 3p microscope, including the voxel acquisition conditions and in particular the optimal choice of the repetition rate, PSF, pulse duration and pulse energy of the excitation laser allow to approach this fundamental depth limit within the linear and non-linear constraints of laser exposure and at adequate spatiotemporal resolutions for cellular and subcellular Ca^2+^ imaging, within moderately sized FOVs. Recording from different cortical and sub-cortical regions within the mouse brain, we demonstrated that our method allows for *in vivo* recording of neuronal Ca^2+^ activity at up to ∼2.1 mm tissue depth within a FOV of ∼230 × 230 µm at ∼ 8.6 Hz.

While we showed that reaching such depths is possible without the implementation of any adaptative optics (AO) strategies^50,51,62,63^, it is clear that the excitation light undergoes major aberrations as it traverses through the refractive index inhomogeneities in tissue. Thus, we expect that in the future, a combination of our method with suitable AO techniques would allow for correction of the PSF aberrations and further enhance in-focus fluorescence, enabling higher-SNR recordings and resolution of finer features at large tissue depths. We do not expect AO methods to enable major increases in imaging depth, since the demonstrated AO signal improvements of ∼5–10 times^50,62,63^ translate only to an imaging depth increase of ∼150–250 μm. However, beyond enabling finer feature resolution, AO could also further reduce average power requirements.

## Methods

### Experimental model and animal preparation

All surgical and experimental procedures were approved by the Institutional Animal Care and Use Committee (IACUC) of The Rockefeller University. Mice were obtained from Jackson Laboratory and bred in-house for the specified transgenic lines. In this study, we used mice expressing green fluorescence protein (Thy1-EGFP, JAX 007788) for static imaging experiments, as well as transgenic lines Camk2a-tTA (JAX 007004) × tetO-GCaMP6s (JAX 024742), Camk2a-tTA (JAX 016198) × tetO-GCaMP8s (JAX 037717), and VGAT-Cre (JAX 016198) × GCaMP8m (JAX 037718) for Ca^2+^ activity imaging.

### Cranial window surgery

Cranial window implantation was performed under isoflurane anesthesia (1–1.5 % maintenance at a flow rate of 0.7–0.9 l/min) using procedures adapted from previous studies^46,48,64^. After mice were placed in a stereotaxic frame (RWD Life Science), dexamethasone (0.4 mg/ml, subcutaneous) and meloxicam (0.125 mg/kg, subcutaneous) were administered preoperatively to reduce brain swelling, suppress inflammatory responses and stabilize the blood-brain barrier. Additionally, bupivacaine (0.125 mg ml^−1^) was injected subcutaneously above the skull to provide local anesthesia and protect tissue from excessive neuronal excitation. The scalp was sterilized with betadine and ethanol, then removed via midline incision to expose the underlying skull. The periosteum and connective tissue were carefully cleared using a scalpel and fine forceps. A custom-made stainless-steel head bar was fixed posterior to the occipital bone using cyanoacrylate glue (Loctite) and reinforced with black dental cement (Ortho-Jet, Lang Dental). A craniotomy was performed using a high-speed dental drill (0.5 mm burr) with continuous irrigation to prevent overheating. The size and shape of the craniotomy varied depending on the imaging requirements, ranging from circular (3 mm diameter) for single-hemisphere imaging to D-shaped (7–8 mm) for enhanced bilateral cortical optical access. A single or two bonded sterile glass coverslips (#1 thickness, Warner Instruments; optical adhesive Norland 60) were implanted into the craniotomy and sealed in place using tissue adhesive (Vetbond).

Following surgery, mice received dexamethasone and meloxicam (as previously described) for seven consecutive days, to minimize immune activation and provide analgesia. Additionally, antibiotic-enriched feed (Sulfatrim) was provided to prevent infection and support physical recovery. Mice were housed in their home cages under standard conditions and allowed to recover for at least 2–3 weeks before imaging. Animals were monitored daily for signs of infection or discomfort and were excluded from imaging experiments upon detection of dura damage or compromised brain tissue clarity.

### TrUDI microscope for 3p imaging

A schematic of the optical path of the 3p microscope is shown in **Extended Data Fig. 4**. To meet the pulse energy requirements as well as the optimal repetition rate across the near-IR and IR wavelength ranges, we used a factory-customized tunable OPA system. The laser system comprised an ultrafast diode-pumped ytterbium-doped regenerative amplifier (Yb:KGW, ytterbium-doped potassium gadolinium tungstate, Carbide CB3, Light-Conversion, 80 W, 500 µJ output at 1030 nm, ∼ 210 fs, 160 kHz) as a pump for the hybrid OPA (Orpheus-F, short pulse mode, Light-Conversion). A part of the pump pulse energy was used to generate a white light continuum by focusing it onto a nonlinear crystal and the rest was used to generate second harmonics required for pumping the OPA stages at 515 nm. The white light was used as the seed for the OPAs by tuning the delay and chirp to select the spectrum and bandwidth for the 3pM wavelength. The weak seed was then amplified in two stages using the SHG pump at the OPA crystals in a hybrid non-collinear and collinear fashion. The idler wave, required for 3p imaging, was then separated from the seed wavelength outputs, and dispersion was pre-compensated to achieve the shortest pulse duration based on the requirement for the downstream optical components. We tuned the OPA to our wavelength of interest, maximizing output power and bandwidth at 1300 nm, resulting in a pulse energy >10 μJ with ∼50 fs compressed pulse duration at the laser output at 160 kHz repetition rate.

For pulse compression and dispersion compensation for the downstream optical components, we used a prism compressor using a pair of Brewster-angle prisms (SF10) placed on an adjustable rail to optimize the pulse duration for 1300 nm. The final pulse duration was optimized at the sample location by collimating the light out of the objective lens followed by free-space coupling to an autocorrelator (APE pulseCheck). At 1300 nm, the minimum pulse duration obtained was approximately 50 fs with a Gaussian fit to the intensity autocorrelation, representing a full bandwidth over 85 nm. For power modulation, we used a fast electro-optic modulator (EOM, Conoptics, 360-40), that handled both primary power adjustment and beam blanking at the turning edge of the fast-scanning axis. However, for imaging across a ∼ 6–7 EAL depth range, the dynamic range for power modulation exceeded the contrast provided by the EOM. Therefore, we incorporated a secondary power adjustment module comprised of a motorized half-wave plate and a polarized beamsplitter (Thorlabs). Beam sizes at various locations were adjusted with telescopes consisting of off-the-shelf lenses (Thorlabs).

The laser output was interfaced with the 3p microscope (Sutter MOM-3P) comprising a 3-axis micrometer stage, galvo-galvo scanners and scan optics. We used a 16× objective with a long working distance of 3 mm and NA 0.8 (Nikon), unless otherwise noted. The detection assembly included a primary dichroic mirror (FF735-Di02 LP) for separating excitation and emission light, an emission dichroic at 565 nm for separating green and red signals, an emission fluorescence filter (525/70 BP for green), and photomultiplier tube (PMT, Hamamatsu, H10770-PA40).

Data acquisition and microscope control were handled with a single data acquisition card (vDAQ, MBF) and ScanImage software (version 2022, MBF). The PMT signal was recorded with a 16-bit digitizer at 120 MS/s sampling rate. A clock generation board (Texas Instruments) provided necessary clock signals derived from the laser repetition rate for synchronized and gated sampling in a single-pulse-per-pixel acquisition scheme. The gate width was optimized to integrate the fluorescence signal for best SNR.

### Imaging protocol

Awake mice were head-fixed on a treadmill platform, and the flat glass window was aligned perpendicular to the optical axis using a tip-tilt stage and an alignment tool based on a red laser diode and beam splitter for generating a reference reflection. In a typical recording the image pixel resolution was 1.8 µm with an image size of 128 × 128 pixels, resulting in a FOV of 230 × 230 µm. In ScanImage, we set the temporal fill fraction of the fast axis to 0.9. With single pulse-per-pixel imaging and a 160 kHz laser, this yielded a frame rate of 8.6 Hz for recording neuronal activity from a single plane. For high-resolution recordings, we used a pixel resolution of 0.9 µm and an image size of 256 × 256 while maintaining the same FOV. For z-stack recordings, the axial resolution was 5–20 µm, as noted in the main text and figure legends for individual cases. Depending on the recording depth, laser power to the EOM was adjusted using the secondary power controller. The EOM adjusted the laser power during changes of depth in single-plane and stack recordings and performed beam blanking during galvo mirror turnaround. The overall range of laser power used was ∼0.5–150 mW, measured after the objective lens with a thermal power sensor (Thorlab S405C) before each imaging session. Since water has substantial absorption at 1300 nm, fresh D_2_O was used as the immersion medium for all recordings.

### Experimental comparison of TrUDI with MHz 3pM

To compare the performance of a MHz imaging system with the optimized TrUDI microscope, we modified the sampling scheme of our 160 kHz system to mimic MHz conditions, enabling a more direct comparison between the two modalities as described in the main text. Briefly, this was achieved by slowing down the scanner to spatially oversample the FOV, matching the sampling of an equivalent 1.3 MHz system, and subsequently averaging the signal from 8 successive pulses into a single pixel to maintain a constant pixel size between configurations. This resulted in an 8-fold higher “extrapolated” average power (*P_extrap_*) in the 1.3 MHz configuration compared to the actual average power (*P_actual_*) used. Using this approach, we performed comparative *in vivo* imaging of a mouse expressing GCaMP8m in the hippocampus through the intact cortex. We compared TrUDI with the mimicked 1.3 MHz configuration under two conditions. In the first, the pulse energy at the focus was kept the same (∼3.1 nJ) for both configurations while imaging at 1080 µm. In the second, the average power was kept the same for both configurations (∼44 mW; i.e., *P_actual_* = 44 mW for TrUDI and *P_extrap_* = 44 mW for the mimicked 1.3 MHz configuration) while imaging at a depth of ∼1.1 mm.

Finally, to compare the imaging of TrUDI and a real 1 MHz system in mouse brain at comparable depth, we constructed a secondary 3p setup following the design in one of our previous publications^37^. Briefly, the 1 MHz system used an OPCPA laser (White Dwarf, Class 5) pumped by fiber laser (80 W, 1030 nm, 1 MHz, Coherent Monaco) to generate 1300 nm light. The laser was then coupled to a Sutter microscope after passing through a power adjustment module, a pulse dispersion compensation module, and a scanner module comprised of a relayed resonance-galvo-galvo scanner configuration, followed by a scan lens (f = 110 mm, Thorlabs LSM05) and tube lens (f = 400 mm, Edmund Optics), and the same objective lens used in TrUDI. The detection system was similar to the TrUDI setup. The data acquisition system consisted of a high-speed FPGA-digitizer assembly (NI 7975R, NI 5772-02, PXI 6341, National Instruments) and ScanImage software (version 2018, MBF). Multi-ROI scanning with a 16 kHz resonant scanner and the galvo-galvo pair was configured to match the pixel resolution, achieve a similar FOV size, and provide an imaging speed of 30 Hz. The system was calibrated to have imaging parameters as close as possible to TrUDI, except for the pulse duration, which for OPCPA source is limited to ∼100 fs (i.e., a factor of two longer than in TrUDI). We performed comparative *in vivo* imaging of deep cortical areas in the PPC of mice expressing GCaMP6s (Camk2a-tTA× tetO-GCaMP6s) on both microscopes (at nearly identical locations) (**Fig. 2a-b**). In TrUDI, we used 7.5 mW average power (∼3 nJ pulse energy at the focus), while for the 1 MHz microscope, we used 193 mW power. Note that for all pulse energy calculations under the experimental conditions described here or elsewhere we assume a depth-dependent exponential loss using an EAL of 300 µm and ignore losses due to the immersion medium or sample surfaces.

### Simulation of three-photon fluorescence signal attenuation at high excitation NA

To investigate how excitation NA and tissue scattering affect 3p fluorescence signal generation, we performed Monte Carlo simulations that modeled ballistic photon propagation followed by nonlinear 3p signal estimation. The simulation assumes a homogeneous scattering medium with an EAL of 300 µm at an excitation wavelength of 1300 nm. The optical system simulated included a paraxial objective lens with a focal length of 12.5 mm (equivalent to focal length of our 16× Nikon objective) and an immersion medium with refractive index 1.33. The initial positions of the photons were drawn from a Gaussian distribution at the back focal plane (BFP), width Gaussian widths chosen to result in a range of excitation NAs (0.2–1.0). Photon trajectories were simulated using a custom Monte Carlo model that launches photons from the BFP and propagates them through the lens and into the tissue. Each photon is directed such that it would reach the geometric focal plane in the absence of scattering. Within the scattering tissue the photons are assigned a travel distance drawn from an exponential distribution characterized by the EAL and for a given focal depth. Photons that reached the focal plane without being scattered were accumulated as ballistic photons and thus the fraction of ballistic photons was computed for each depth and excitation NA.

To quantify the 3p fluorescence signal, we calculated two scenarios: one that ignores angular-dependent marginal ray attenuation (thus the loss of photons only depends on the focal depth), and one that incorporates scattering-induced losses of the marginal rays using the Monte Carlo results. In the former case, the pulse energy at the sample is compensated for the depth-dependent exponential loss, making the 3p signal independent of focal depth, but increasing with the excitation NA as expected. In the latter case, identical pulse energy compensation as in the former case was also applied for the given depth and the fraction of ballistic photons from the Monte Carlo simulation was used to account for the loss of marginal rays, which becomes significant at high excitation NAs. In both cases, the resulting nonlinear 3p fluorescence signals were normalized to the maximum signal for the case without marginal ray attenuation for comparison.

### Data analysis pipeline

The raw tiff data obtained from ScanImage were deinterleaved to extract channels of interest, followed by residual line phase offset correction due to the bidirectional scan along the fast axis. For long recordings, motion correction was performed using the NoRMCorre library^65^. We used the following parameters for the rigid motion correction (see NoRMCorre documentation): upsampling factor 50, maximum shift 15 pixels, initial batch size 200, and bin width 200. Other parameters were kept at their defaults. Example of motion correction metrics are shown in **Extended Data Fig. 5** for two datasets at imaging depths 730 µm and 1840 µm (corresponding to the single plane Ca^2+^ imaging data in **Fig. 4**), indicating good registration. For 3D volume reconstruction of functional imaging data, mean projections were calculated from single plane data with axial separation 5–20 μm. Spatial components of neuron region of interests (ROIs) were extracted using Cellpose in MATLAB.^66^ The individual planes were then combined to single volumes (**Fig. 4b** and **Fig. 5b**). For structural imaging with GFP mice, 3D volumes were reconstructed from the raw data to keep the fine structures of the neurons as shown in **Fig. 3**.

To extract neuron spatial footprints and deconvolved activity traces, we used the CNMF package CaImAn^61^ that performs spatio-temporal filtering and constrained non-negative matrix factorization to find components. We applied a rolling mean filter of window size 5 frames on the motion- and phase-corrected datasets before running it through CaImAn pipeline. Key CaImAn settings included (see CaImAn documentation): initialization method ‘greedy’, search method ‘ellipse’, deconvolution ‘constrained FOOPSI-SPGL1’, autoregression ‘decay’, estimated ROI width 8 pixels, marge threshold 0.9, fudge factor 0.99, frame rate 8.6, min. SNR 1.5, temporal iterations 5. Other parameters were kept at their defaults. The components were manually classified based on the ROI refinement feature of the CaImAn-based EZcalcium^67^ toolbox with parameters mean width (5–10 pixel), roundness (>0.8), oblongness (<2) and sorted by activity (z-score). The results are shown in **Fig. 4**, **Fig. 5, Extended Data Fig. 6,** and **Extended Data Fig. 7**. For all time-projected images shown, the contrast or color saturation was adjusted to aid visualization.

## Data availability

All source data supporting the findings of the study are provided with the paper. The raw data are available from the corresponding author upon reasonable request.

## Code availability

Neuronal segmentations based on anatomical or spatio-temporal feature, and motion-corrections are implemented in MATLAB using open-source packages and are publicly available. All custom codes used for data analysis in this work are provided as compressed archive and available with the paper as **Supplementary File 1**.

## Acknowledgements

Research reported in this publication was supported by the National Institute of Neurological Disorders and Stroke of the National Institutes of Health under award numbers 5U01NS115530 (A.V.), 5U01NS126057 (A.V.), 1RF1NS113251 (A.V.), 1RF1NS110501 (A.V.), by the Kavli Foundation through the Kavli Neural System Institute (A.V.), by a Pershing Square award through the William Ackman and Neri Oxman Innovation Fund in Neurodegenerative Diseases (A.V.) and the Torsten N. Wiesel Fellowship in Mind, Brain and Behavior at the Rockefeller University (K.S.).

## Author Contributions

K.S. contributed to the conceptualization of the project, built and characterized the imaging system, implemented the data acquisition system, programmed experimental control and analysis software, performed experiments, analyzed data, and wrote the manuscript. K.B., F.L. and A.F. performed cranial window surgeries. A. B. and J. L. provided additional transgenic animals with cranial windows. A.V. conceived and led the project, conceptualized, designed and guided the implementation of the imaging system and data analysis and wrote the manuscript.

