## Supplementary Information for "Cellular-resolution Three-photon Ultra-Deep Imaging (TrUDI) of neuronal population activity at 2 mm depth in the mammalian brain"

#### Supplementary Notes

##### *Supplementary Note 1*

*Signal-to-Background Ratio in 3p imaging*

In multiphoton microscopy, the generation of the fluorescence signal relies on the $n$-photon absorption process, which depends on the $n$-th power of the beam intensity, the multiphoton absorption cross-section, $\sigma^{(n)}$, the concentration of the fluorophores, $C$, and the excitation volume under consideration. For a Gaussian pulsed beam propagating through a transparent medium, the intensity can be written as:

$$I(r,z,t) = I_0 \left(\frac{w_0}{w_z}\right)^2 \exp\left(-\frac{2r^2}{w_z^2}\right) \exp\left(-\frac{2t^2}{\tau^2}\right) \tag{1}$$

where $I_0$ is the peak intensity, which is proportional to $E_0/w_0^2\tau$, $E_0 = P_0/f_R$ is the pulse energy, and $P_0$ is the average power at the surface defined by the depth $z = 0$, $w_0$ is the focal spot size (1/e[2] radius and given by $\frac{\lambda}{\pi\, NA}$, where $NA$ is the effective numerical aperture), $w_z$ is the beam size at an arbitrary location $z$, $\tau$ is the pulse duration (FWHM), $f_R$ is the laser repetition rate. In tissue samples, however, the intensity of the beam attenuates exponentially as a function of depth due to absorption and scattering, which is characterized by an effective attenuation length (EAL), $l_{np}$. Therefore, the modified intensity can be written as,

$$I(r,z,t) = I_0 \left(\frac{w_0}{w_z}\right)^2 \exp\left(-\frac{2r^2}{w_z^2}\right) \exp\left(-\frac{2t^2}{\tau^2}\right) \exp\left(-\frac{z}{l_{np}}\right) \tag{2}$$

The fluorescence signal can be written as,

$$F(r,z,t) \propto \frac{1}{n} \sigma^{(n)} g^{(n)}\, C\, I^n(r,z,t) \left(\frac{\lambda}{hc_0}\right)^n \tag{3}$$

where, $g^{(n)}$ denotes the $n$-th order temporal coherence of the pulse, $\lambda$ is the excitation wavelength, $h$ is Planck's constant, and $c_0$ is the speed of light in vacuum. The fluorescence signal generated from a thin slice of tissue of thickness $dz$ at a depth of $z$ (with effective volume, $dV = \pi w_z^2 dz$) due to a single pulse of duration $\tau$ can be obtained from equation (3) via integration over $r$ and $t$, followed by substitution of the expression for $I_0$, as

$$F(z) \propto \frac{1}{n} \sigma^{(n)} g^{(n)} \left(\frac{\lambda}{hc_0}\right)^n C \frac{E_0^n \exp\left(-\frac{nz}{l_{np}}\right)}{\tau^n w_z^{2n}} dV\tau \tag{4}$$

For 3pM, we can simply write

$$F(z) \propto \frac{E_0^3 \exp\left(-\frac{3z}{l_{3p}}\right)}{\tau^2 w_z^6} dV \qquad (5)$$

which can also be expressed in terms of average power $P_0$ as,

$$F(z) \propto \frac{P_0^3 \exp\left(-\frac{3z}{l_{3p}}\right)}{\tau^2 f_R^3 w_z^6} dV \qquad (6)$$

and thus, for a given average power, the fluorescence signal increases nonlinearly with decreasing laser repetition rate.

The focal fluorescence signal generated per pulse from a focal volume of $2\pi w_0^2 z_R$ is given by

$$F_f \propto \frac{E_0^3 \exp\left(-\frac{3z_f}{l_{3p}}\right)}{\tau^2 w_0^4} z_R \qquad (7)$$

and the out-of-focus background fluorescence can be written as

$$F_{oof} \propto \left[\int_0^\infty \frac{E_0^3 \exp\left(-\frac{3z}{l_{3p}}\right)}{\tau^2 w_z^4} dz\right] - F_f \qquad (8)$$

where $z_R$ is the Rayleigh range of the Gaussian beam at the focus $z_f$. Finally, we can write the expression for signal-to-background ratio (SBR) as,

$$1/SBR = F_{oof}/F_f = \left[\int_0^\infty \frac{\mathrm{z}_\mathrm{R}^3 \exp\left(-\frac{3(z-z_f)}{l_{3p}}\right)}{\left[z_R^2 + \left(z - z_f\right)^2\right]^2} dz\right] - 1 \qquad (9)$$

Using the SBR calculated for a range of focal depths $z_f$, the depth limit $z_{DL}$ can be obtained by solving for SBR at unity as shown in **Fig. 1a**.

***Supplementary Note 2***

*Single-pulse-per-pixel in 3p imaging*

To achieve the ultra-deep tissue imaging, we utilized optimal excitation by employing a single-pulse-per-pixel scheme. For the non-linear excitation, the fluorescence signal at the focus from $p$ pulses each with energy $E_0$, can be obtained from equation (7) as

$$F_{f,p} \propto p \; \frac{E_0^3 \exp\left(-\frac{3z_f}{l_{3p}}\right)}{\tau^2 w_0^4} z_R \tag{10}$$

Using the same average power $P_0$, but with a single high-energy pulse with energy $pE_0$, the fluorescence signal at the focus becomes

$$F_{f,1h} \propto \frac{(pE_0)^3 \exp\left(-\frac{3z_f}{l_{3p}}\right)}{\tau^2 w_0^4} z_R \tag{11}$$

Therefore, $F_{f,1h} = p^2 F_{f,p}$, i.e., for same average power the signal is maximized with single-pulse excitation, which is the minimum requirement in order to form a pixel and consequently maximizing the SNR under shot-noise-limited condition. For a fixed average power, the repetition rate of the laser must be scaled down by the same gain factor, *i.e.,* $f_{R2} = f_{R1}/p^2$. Thus, the nonlinear gain suggests that it is crucial to use a laser with low repetition rate and high pulse energy in a single-pulse-per-pixel fashion in order to minimize average power while maximizing signal for deep tissue imaging.

***Supplementary Note 3***

*Mean photon number as the detected 3p fluorescence signal for a given pulse energy at the focus*

In order to estimate the number of photons ($N$) per pulse arriving at the PMT, we can use equations (3) and (7) to write the theoretical estimates, assuming the photons are generated primarily at the focus, as

$$N = \frac{1}{3}\eta g^{(3)} \sigma^{(3)} C N_A \phi^3 V \tau \tag{12}$$

where $\phi = I_0\lambda/hc_0$ is the photon flux (photons per unit area per unit time) and $I_0$ is the intensity. $\eta$ is the overall photon generation and collection efficiency of the system, $g^{(3)}$ denotes the 3rd order temporal coherence of the pulse, $\sigma^{(3)}$ is the 3-photon absorption cross-section of the fluorophores. $N_A$ represents Avogadro's number and $C$ is the molar concentration of the fluorophore at the focal volume $V = 2\pi w_0^2 z_R$ within the Rayleigh range $z_R$. Within the focal volume, the photon flux can be approximated by

$$\phi = \frac{\lambda E_0}{hc_0\pi w_0^2\tau} \quad (13)$$

Where $E_0$ is the pulse energy at the focus, $\lambda$ is the excitation wavelength, $h$ is Planck's constant, $c_0$ is the speed of light in vacuum, $w_0$ is the focal spot size (1/e$^2$ radius and given by $\frac{\lambda}{\pi NA}$, where $NA$ is the effective numerical aperture), $\tau$ is the pulse duration (FWHM). Since the fluorescence photons arriving at the detector follow Poisson statistics, equation (12) represents the mean photon number in a given time interval. As an example, with a pulse energy $E_0$ of 2.25 nJ at 1300 nm, $NA$ = 0.25, $\tau$ = 50 fs (FWHM), 3-photon cross-section $\sigma^{(3)}$ = $19 \times 10^{-95}$ m$^2$s$^2$photon$^{-2}$, $g^{(3)}$ = 0.51, refractive index 1.33, overall efficiency $\eta$ = 0.05, and molar concentration of fluorophore $C$ = 10 μM, we have $N$ ~ 3 for the average number of photons arriving at the PMT.

*Theoretical estimation of the system noise and SNR for an average of N fluorescence photons*

The $N$ photons arriving at the detector produce photoelectrons based on the quantum efficiency of the PMT cathode, which then undergo multistage amplification at the dynode. Due to the random nature of the gain, the final current output at the PMT contains extra noise in addition to the photon shot noise. Since the number of photons arriving at the detector is low, the PMT will generate a short pulse with a finite width $\delta T$. As the fluorescence photons are Poisson-distributed, the PMT response time to the $N$ photon signal is be a convolution of the PMT's single-photon response and the fluorescence lifetime of the fluorophore molecule. The anode current ($I_a$) signal is then given as the product of optical power, mean gain and cathode radiant sensitivity of the PMT as follows:

$$I_a = \frac{GS_c hc_0}{\lambda_{em}} \frac{1}{\delta T} N = \alpha N \tag{14}$$

where $G$ is the mean gain of the PMT, $S_c$ is the PMT cathode radiant sensitivity (defined as the photocurrent generated at the cathode per unit incident power), $\delta T$ is the PMT response time of $N$ fluorescence photons, and $\lambda_{em}$ is the emission wavelength. The term $\alpha$ encapsulates all other terms except $N$. For our system, typical values of the parameters are: $G = 2 \times 10^6$, $S_c = 189 \times 10^{-3}$ A/W, $\lambda_{em}$ = 520 nm and $\delta T$ = 3.5 ns. Thus, $\alpha = 4.13 \times 10^{-5}$.

The variance of the anode signal can be expressed as,

$$\sigma_{I_a}^2 = \left(\frac{GS_c hc_0}{\lambda_{em}} \frac{1}{\delta T}\right)^2 N + (2eBFG)\left(\frac{GS_c hc_0}{\lambda_{em}} \frac{1}{\delta T}\right) N = \alpha(\alpha + \beta) N \tag{15}$$

where $\beta = 2eBFG$ is a PMT noise contributing factor, where $e$ is the charge of an electron, and $B$ is the bandwidth of the system which, in absence of other bandwidth-limiting factors, can be approximated as $B \approx 0.35/T_r$, with rise-time $T_r \approx \delta T/2$. $F$ is the noise figure in the multi-stage process (typically close to one and can be neglected). Thus, for our system, $\beta = 1.28 \times 10^{-4}$. The signal-to-noise ratio (SNR) is then given by:

$$SNR = \frac{I_a}{\sigma} = \sqrt{\frac{\alpha}{\alpha + \beta}} \sqrt{N} \tag{16}$$

In order to express the PMT noise in terms of image pixel intensity $Y_{img}$, we can convert the anode current to the measured voltage at the analog input of the data acquisition system followed by the digitization of the signal. For our system, the relationship between anode current and digital counts is:

$$Y_{img} = \kappa I_a = \kappa\alpha N \tag{17}$$

where, $\kappa$ is the conversion factor from PMT current to pixel digital counts, given by $R/r_b$, where $r_b$ is the digitizer resolution in volts per count (30.5 µV/count) and R is the equivalent load resistance at the analog input (50 Ohm). Thus, $Y_{img} \approx 67.7\ N$, and we can rewrite the equations (15) and (16) in terms of image pixel intensity.

$$\sigma_{img}^2 = \kappa(\alpha + \beta)Y_{img} \tag{18}$$

and

$$SNR_{img} = \sqrt{\frac{1}{\kappa(\alpha + \beta)}}\sqrt{Y_{img}} = \frac{1}{\gamma}\sqrt{Y_{img}} \tag{19}$$

where $\gamma$ is a constant. Using the typical values of the parameters of our system, we have theoretical value of $\gamma \approx 16.6$.

*SNR of the baseline fluorescence trace of neurons of at the SBR limited condition*

Equation (16) is valid when out-of-focus background is ignored ($SBR \gg 1$). In the limit of SBR = 1, however, this must be incorporate the noise contribution of the background photons. If $N_B$ represent the number of background photon per pulse, we can rewrite the SBR dependent SNR as:

$$SNR = \sqrt{\frac{\alpha}{\alpha + \beta}}\sqrt{\frac{N^2}{N + N_B}} \tag{20}$$

Thus, by rearranging the terms and substituting the definition of $SBR = N/N_B$, we obtained:

$$SNR = \sqrt{\frac{\alpha}{\alpha + \beta}}\sqrt{\frac{1}{1 + 1/SBR}}\sqrt{N} \tag{21}$$

For single-pulse-per-pixel imaging scheme, equation (21) represents the single-pixel baseline SNR ($SNR_{base,\ pix}$) as defined in the main text. For neuronal imaging, the overall signal from the neuron, and thus the SNR, will improve with number of pixels sampled per neuron. If $M_{pix}$ is the number of pixels per neurons per frame, then the variance is $\sigma^2_{I_a,M_{pix}} = \alpha(\alpha+\beta)(N+N_B)M_{pix}$ and the SNR is $SNR_{M_{pix}} = \sqrt{\frac{\alpha}{\alpha+\beta}}\sqrt{\frac{1}{1+1/SBR}}\sqrt{NM_{pix}}$. For the neuron baseline signal this is represented by baseline the SNR ($SNR_{base}$) as defined in the main text. In **Fig. 1b**, the $SNR_{base}$ of a neuron is shown as functions of mean number of detected photons per pixel ($N$) for different neuron sizes for both in absence of background as well as at the SBR limit of 1. In **Extended Data Fig. 1a-b**, the required $SNR_{base}$ threshold and the corresponding variance of the PMT output current per pixel are shown as functions of mean number of detected photons per pixel ($N$) for the desired $SNR_{base}$ = 3 per neuron. Furthermore, for the given desired baseline $SNR_{base}$, the required number of photons per neuron per second i.e., the baseline photon detection rate ($F_0$) for a neuron can be obtained as follows:

$$F_0 = (SNR_{base})^2\left(\frac{\alpha+\beta}{\alpha}\right)\left(1+\frac{1}{SBR}\right)f_R \qquad (22)$$

where, $f_R$ is the imaging frame rate. Since we assumed $SNR_{base}$ = 3 for the neuron, therefore at SBR-limited depth (SBR = 1) at a imaging frame rate of 8.6 Hz, this leads to baseline fluorescence detection rate of ~635 photon per second per neurons given in the main text.

*Experimental evaluation of the system noise and SNR under in-vivo imaging conditions*

We estimated the signal (mean pixel count) and the corresponding noise (variance and SNR of the pixel count) from the experimental measurement. Using the optimal PSF size and require pulse energy, we performed *in vivo* imaging with the typical condition, i.e., with a ~2 nJ pulse energy at the focus, a field of view (FOV) of 230 × 230 µm$^2$, pixel size of 1.8 × 1.8 µm$^2$ at 8.6 Hz frame rate and using the single-pulse-per-pixel data acquisition scheme mentioned in the main text. As shown in **Extended Data Fig. 2**, the measured variance of the pixel count is linear with the mean pixel count, with a slope of ~280. Furthermore, the measured pixel SNR fits the power law given in equation (19) with exponent ~0.5 and pre-factor of ~0.06, as shown in **Extended Data Fig. 2b**. From either the slope of the plot in **Extended Data Fig. 2a** or the pre-factor in **Extended Data Fig. 2b**, we find that the experimental value of $\gamma$ is ~ 16.7, which agrees with the theoretical model above very well.

***Supplementary Note 4***

*Discriminability index as the indicator of neuronal activity signal-to-noise ratio*

In order to determine the minimum requirement to identify neurons' $Ca^{2+}$ activity systematically and reliably from their baseline fluorescence we followed the treatment provide in Ref. 2 in terms of discriminability index, which, in the absence of any out-of-focus fluorescence background, is defined as:

$$d' = \frac{\Delta F}{F}\sqrt{\frac{F_0 \tau_{Ca}}{2}} \tag{23}$$

where $\frac{\Delta F}{F}$ is the relative fluorescence change induced by one action potential (AP), with $F_0$ the baseline fluorescence photon detection rate and $\tau_{Ca}$ is the half-life of the $Ca^{2+}$ decay. In the case where the background generated from the out-of-focus fluorescence is no longer negligible, such as in our aim of imaging at the SBR-limited depth, let us denote $F_{0,bg} = F_0 + B$ and $F_{bg}(t) = F(t) + B$ as the observed baseline fluorescence and observed fluorescence at any arbitrary time point ($t$), respectively, in presence of background ($B$). In equation (23) we can replace the fluorescence-dependent terms with new quantities $\left(\frac{\Delta F}{F}\right)_{bg}$ and $F_{0,bg}$, respectively. The former can be written as:

$$\left(\frac{\Delta F}{F}\right)_{bg} = \frac{F_{bg}(t) - F_{0,bg}}{F_{0,bg}} = \frac{F(t) - F_0}{F_0}\frac{F_0}{F_{0,bg}} = \frac{\Delta F}{F}\frac{F_0}{F_0 + B} \tag{24}$$

Substituting the background-depended terms in equation (23) we can rewrite it as:

$$d' = \frac{\Delta F}{F}\frac{F_0}{F_0 + B}\sqrt{\frac{(F_0 + B)\tau_{Ca}}{2}} \tag{25}$$

Using the definition $SBR = F_0/B$ for the baseline, this can be simplified to:

$$d' = \sqrt{\frac{F_0}{F_0 + B}}\frac{\Delta F}{F}\sqrt{\frac{F_0\tau_{Ca}}{2}} = \sqrt{\frac{1}{1 + 1/SBR}}\frac{\Delta F}{F}\sqrt{\frac{F_0\tau_{Ca}}{2}} \tag{26}$$

Finally, we note that if multiple APs occur within $\tau_{Ca}$, the observed $\Delta F$ and therefore $\frac{\Delta F}{F}$ will scale linearly with the number of APs ($n_{AP}$) for small $n_{AP}$[1]. Thus, the AP-dependent discriminability index can be written as:

$$d'(n_{AP}) = \sqrt{\frac{1}{1 + 1/SBR}}n_{AP}\frac{\Delta F}{F}\sqrt{\frac{F_0\tau_{Ca}}{2}} \tag{27}$$

Equations (26) and (27) are discussed in the main text.

*Detection accuracy metric for the calcium activity of neuron from the discriminability index*

The discriminability index presented in the above section is obtained from the log-likelihood ratio of obtaining a given signal in the presence of spike events and in the absence of spike events (i.e., baseline)[2]. In presence of sufficient signal, the log-likelihood ratio ($\mathcal{L}$) is assumed to be Gaussian distributed with means $\mu_L^{(1)}$ and $\mu_L^{(0)}$ in presence and absence of a spike event, respectively, and are assumed to have the same standard deviations in the case with spike and without spike (i.e., $\sigma_L^{(1)} = \sigma_L^{(0)} = \sigma_L$). Using the definition of discriminability index from signal detection theory:

$$d' = \frac{\mu_L^{(1)} - \mu_L^{(0)}}{\sigma_L} \qquad (28)$$

The true positive rate (TPR) and the false positive rate (FPR) are given by the cumulative distribution function $\Phi(x)$ of the Gaussian distribution as follows:

$$TPR = 1 - \Phi\left(\frac{\xi - \mu_L^{(1)}}{\sigma_L}\right);\ FPR = 1 - \Phi\left(\frac{\xi - \mu_L^{(0)}}{\sigma_L}\right) \qquad (29)$$

where $\xi$ is determined by the decision threshold for identifying a spike events (i.e., if $\mathcal{L} > \xi$, then spike events occurred, and vice versa). For the log-likelihood ratio the "unbiased criterion" can be obtained with $\xi = 0$, and additionally it has been shown that the distribution of the log-likelihood ratio is symmetric around the criterion, yielding $\mu_L^{(1)} = -\mu_L^{(0)}$.[2] Using this relation in equation (28), we have $\frac{\mu_L^{(1)}}{\sigma_L} = -\frac{\mu_L^{(0)}}{\sigma_L} = \frac{d'}{2}$. Furthermore, we can simplify equation (29) by using these relations as follows.

$$TPR = \Phi\left(\frac{\mu_L^{(1)}}{\sigma_L}\right) = \Phi\left(\frac{d'}{2}\right);\quad FPR = \Phi\left(\frac{\mu_L^{(0)}}{\sigma_L}\right) = \Phi\left(-\frac{d'}{2}\right) \qquad (30)$$

We used the identity $\Phi(-x) = 1 - \Phi(x)$. These relations of TPR and FPR with respect to $d'$ have been used in the main text to obtain $Ca^{2+}$ activity detection TPR >95.5% and FPR <4.5% when $d'$ values are 3.4, 6.8 and 10.1 for 1, 2, and 3 APs respectively. Since $d'$ depends on the baseline fluorescence rate ($F_0$), plots for TPR and FPR as shown in **Extended Data Fig. 1c-d** as a function of $F_0$ for a neuron.

### Extended Data Figures

**Extended Data Fig. 1** **Baseline signal-to-noise ratio (SNR) thresholds, neuron activity detection metrics, and NA-dependent fluorescence signals generated in three-photon (3p) microscopy corresponding to the optimal parameter choices shown in Fig. 1**

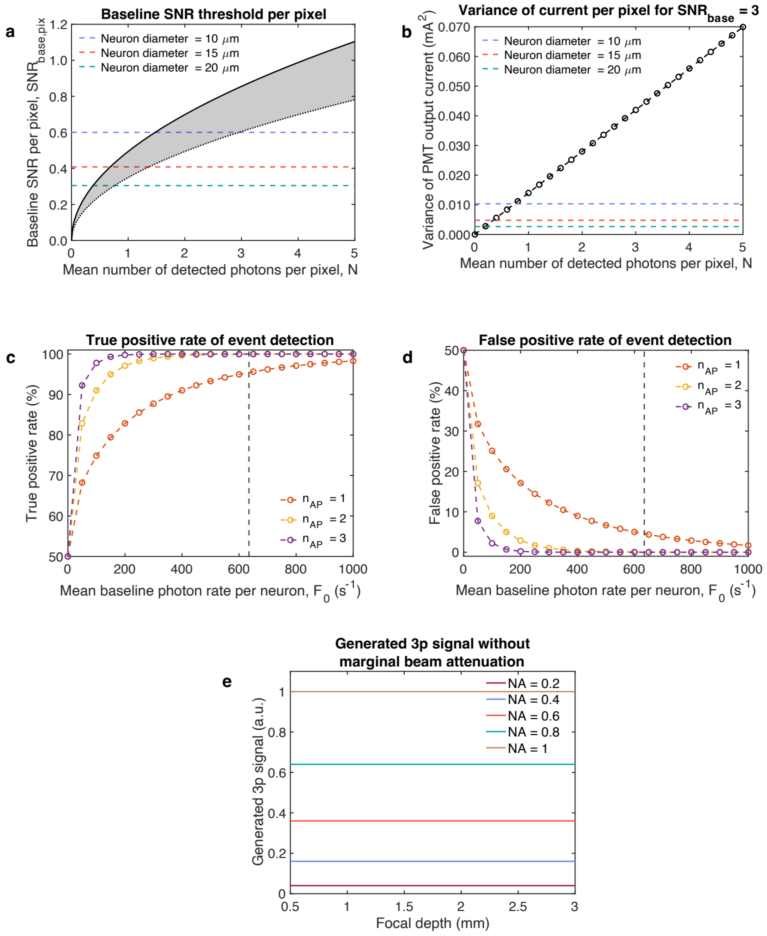


**Extended Data Fig. 1:**
**Baseline SNR thresholds, neuron activity detection metrics, and NA-dependent fluorescence signals generated in 3p microscopy corresponding to the optimal parameter choices shown in Fig. 1.**
**(a-b)** Baseline SNR thresholds and variance of the PMT output current per pixel for the criterion shown in **Fig. 1b**: (a) Baseline SNR per pixel ($SNR_{base,\,pix}$) as functions of the mean number of detected photons per pixel at the PMT. The plot for $SNR_{base,\,pix}$ is shown for both conditions where background is negligible (i.e., SBR >> 1, upper solid line of the shaded region) as well as at SBR-limited depth (i.e., SBR = 1, lower dotted line of the shaded region). The horizontal dashed lines represent minimum threshold for baseline SNR per pixel to achieve baseline SNR of 3 per neuron for three different neuron sizes with diameters of 10, 15, and 20 μm. (b) Variance of PMT output current as functions of the mean number of detected photons per pixel at the PMT. The horizontal dashed lines represent the threshold to achieve baseline SNR of 3 per neuron for three different neuron sizes with diameters of 10, 15, and 20 μm. For (a-b), the theoretical model and the parameters are provided in **Supplementary Note 3**.

**(c–d)** Detection accuracy indicators of an underlying neuronal activity event for the discriminability index shown in **Fig. 1c** at the SBR-limited depth (SBR = 1): (c) True positive rate (TPR) and (d) false positive rate (FPR) are plotted as a function of mean baseline photon rate per neuron (per second). In (c-d), the vertical dashed lines represent the mean baseline photon rate per neuron sampled with 25 pixels at a rate of 8.6 Hz in order to achieve a minimum baseline SNR threshold of 3 per neuron.
(**e**) Generated 3p fluorescence signal as a function of focal depth for different excitation NAs under the assumption that scattering-related photon loss depends solely on depth, and attenuation of marginal rays is ignored, unlike as shown in **Fig. 1d** (which demonstrate the additional attenuation via Monte-Carlo simulation). Note: Pulse energy at the sample is compensated for the depth-dependent exponential loss, making the 3p signal independent of focal depth but increasing with the excitation NA.

**Extended Data Fig. 2** Experimental evaluation of system noise and signal-to-noise ratio (SNR) and comparison with theoretical SNR model in three-photon microscopy

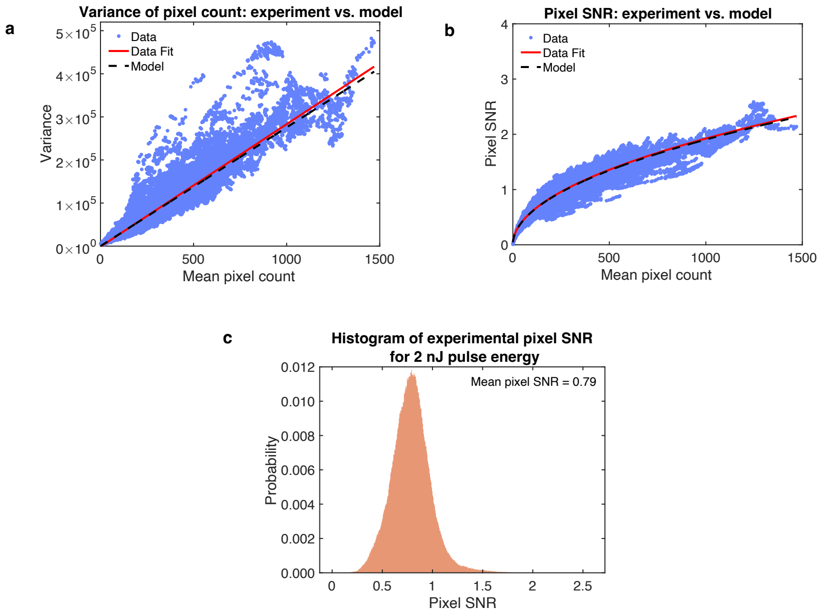


**Extended Data Fig. 2:**

**Experimental evaluation of system noise and SNR and comparison with theoretical SNR model in three-photon microscopy.**

(**a**) Variance of pixel count as a function of mean pixel count from raw image data. The slope of the linear fit (red solid line) is ~280, and the fit agrees well with the theoretical model for variance (green dashed line) (**Supplementary Note 3**).

(**b**) Pixel SNR as a function of mean pixel count from raw image data. A power-law, $y = Ax^B$, fits the data well, with coefficients $A = 0.06$ and $B = 0.5$, in good agreement with the theoretical SNR model (green dashed line) (**Supplementary Note 3**).

(**c**) Histogram of experimental pixel SNR over the FOV across multiple 100-frame windows.

The experimental data were collected under the typical conditions used in our imaging system, i.e., FOV of 230 × 230 μm$^2$; pixel size of 1.8 × 1.8 μm$^2$ at an 8.6 Hz frame rate. We used ~2 nJ pulse energy at the focus using the single-pulse-per-pixel data acquisition scheme. To estimate pixel statistics, we used a 100-frame window. The theoretical model and the parameters are given in **Supplementary Note 3**.

**Extended Data Fig. 3** **Laser-induced non-linear damage thresholds at the tissue surface for hypothetical extended depth reach in three-photon microscopy**

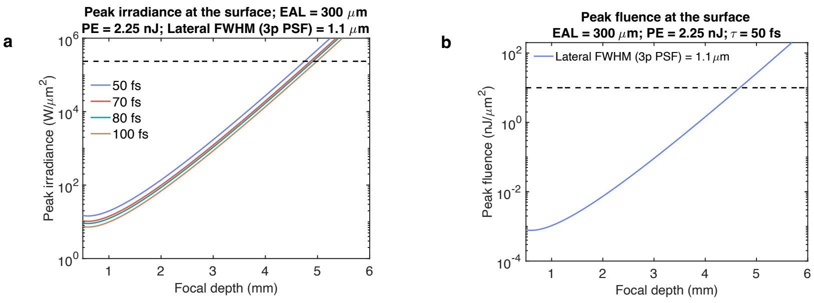


#### Extended Data Fig. 3:

#### Laser-induced non-linear damage thresholds at the tissue surface for hypothetical extended depth reach in three-photon microscopy.

(**a**) Extended plots relating to **Fig. 1g** for focal depth ranges extended to a hypothetical 6 mm depth. The peak irradiance at the surface as a function of focal depth for a pulse energy (PE) of 2.25 nJ, the PSF size as shown in **Fig. 1e**, and an assumed EAL of 300 μm at 1300 nm. Different colored plots correspond to for different pulse durations. The horizontal dashed line corresponds to the peak irradiance limit of ~$2.35 \times 10^5$ W/μm$^2$ for the safe exposure[3].

(**b**) Extended plots of **Fig. 1h** for focal depth ranges extended to a hypothetical 6 mm depth. The peak fluence at the surface as a function of focal depth for the PSF size shown in Fig. 1**e**, PE of 2.25 nJ, and pulse duration of 50 fs (FWHM) assuming an EAL of 300 μm at 1300 nm. The horizontal dashed line corresponds to the peak fluence limit of 10 nJ/μm$^2$ for the safe exposure.[3]

Extended Data Fig. 4 Schematic of the optical setup for Three-photon Ultra-Deep Imaging (TrUDI)

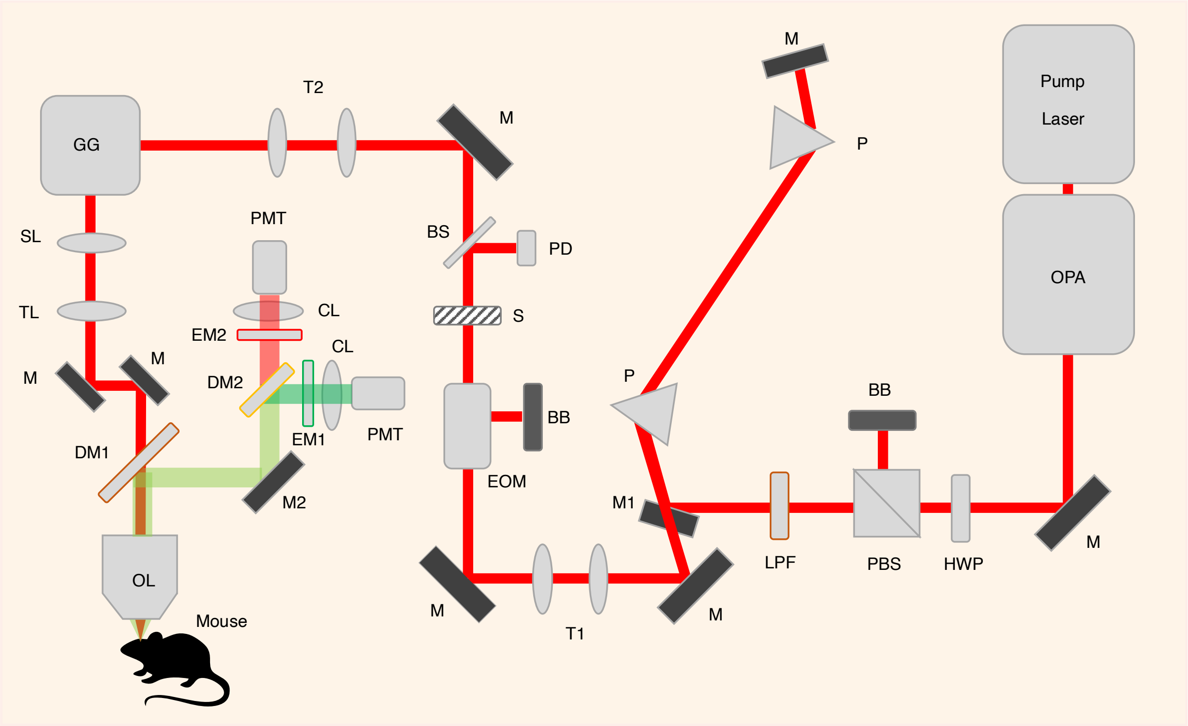


**Extended Data Fig. 4:**
**Schematic of the optical setup for Three-photon Ultra-Deep Imaging (TrUDI).**
TrUDI microscope beam path schematic with a galvo-galvo scanner, a power control module and a dispersion compensation module. Pump Laser (Light-Conversion, Carbide CB3 80W); OPA (Light-Conversion, Orpheus-F); M, mirror (Flat silver mirror Thorlabs PF10-03-P01 or similar); HWP, half-waveplate (Thorlabs AHWP10M-1600); PBS, polarizing beam-splitter (Thorlabs PBS124); BB, beam-block; LPF, 900 nm long-pass filter (Thorlabs FELH0900); M1, D-shaped mirror for beam pick-up (Thorlabs PFD10-03-P01); P, prism-pair for dispersion compensation (Newport 10SF10); T1–T2, telescope to resize laser beam; EOM, electro-optical modulator for fast power adjustment (Conoptics Model 160-40); S, laser shutter (Thorlabs SH05/M); BS, beam sampler (Thorlabs BSF05-C); PD, photodiode (Thorlabs PDA20CS2); GG, galvo-galvo scanner, 5 mm (Novanta, 6215H XY scanner); SL, scan lens f = 50 mm (Thorlabs SL50-3P); TL, tube lens, f = 200 mm (Thorlabs TL200-3P); Sutter MOM-3P detection optics (DM1, Semrock FF735-Di02-25x36; M2, elliptical curved mirror; DM2, Chroma 565DCXR; EM1, Chroma ET525/70m; EM2, Chroma ET605/70m; CL, condenser collection lenses; PMT, Hamamatsu H10770PA-40); OL, objective lens (Nikon 16×, 0.8 NA, 3 mm WD, CFI75LWD16xW2).

**Extended Data Fig. 5** **Estimation of motion artifacts and their correction in deep 3p imaging in the mouse brain in the posterior parietal cortex (PPC) and hippocampus (HPC) for the data shown in Fig. 4**

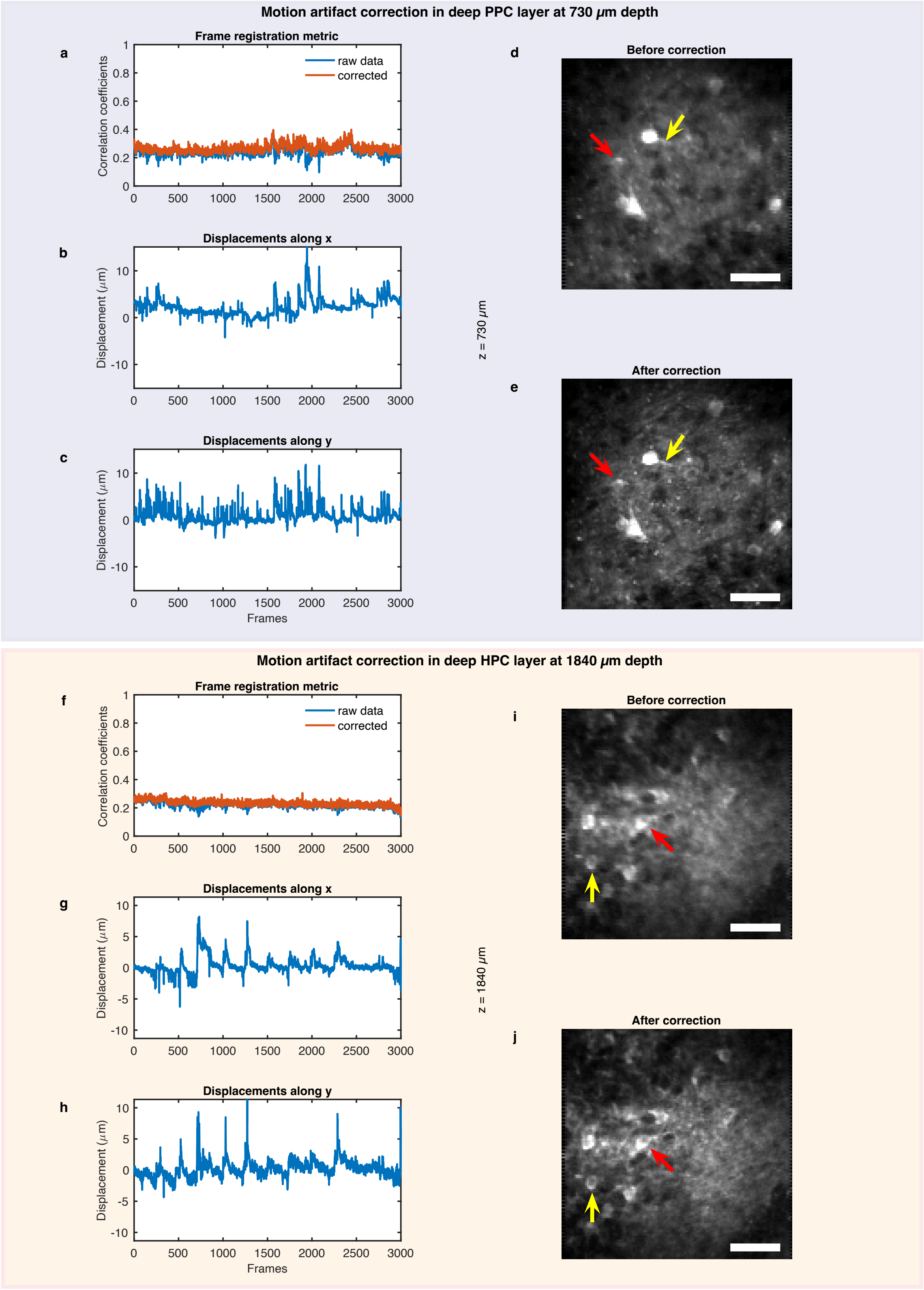

**Extended Data Fig. 5:**
**Estimation of motion artifacts and their correction in deep 3p imaging in the mouse brain on the posterior parietal cortex (PPC) and hippocampus (HPC) for the data shown in Fig. 4.**
(**a-e**) Motion artifact correction in deep PPC layer at 730 µm depth. Correction metrics: (a) Correlation coefficient as frame registration metric for raw and motion-corrected data. (b-c) Detected displacements along x and y in a single plane caused by motion, respectively. (d-e) Mean projection of the single-plane recording before and after motion correction, respectively. The red and the yellow arrows highlights reduced motion blur at two locations. Scale bar, 50 µm.
(**f-j**) Motion artifact correction in a deep HPC layer at 1840 µm depth. Correction metrics: (f) Correlation coefficient as frame registration metric for raw and motion-corrected data. (g-h) Detected displacements along x and y in a single plane caused by motion, respectively. (i-j) Mean projection of the single-plane recording before and after motion correction, respectively. The red and the yellow arrows highlights reduced motion blur at two locations. Scale bar, 50 µm.

**Extended Data Fig. 6** **All segmented neurons and corresponding CaImAn extracted calcium ($Ca^{2+}$) activity traces across depths in posterior parietal cortex (PPC) and hippocampus (HPC) for the data corresponding to Fig. 4**

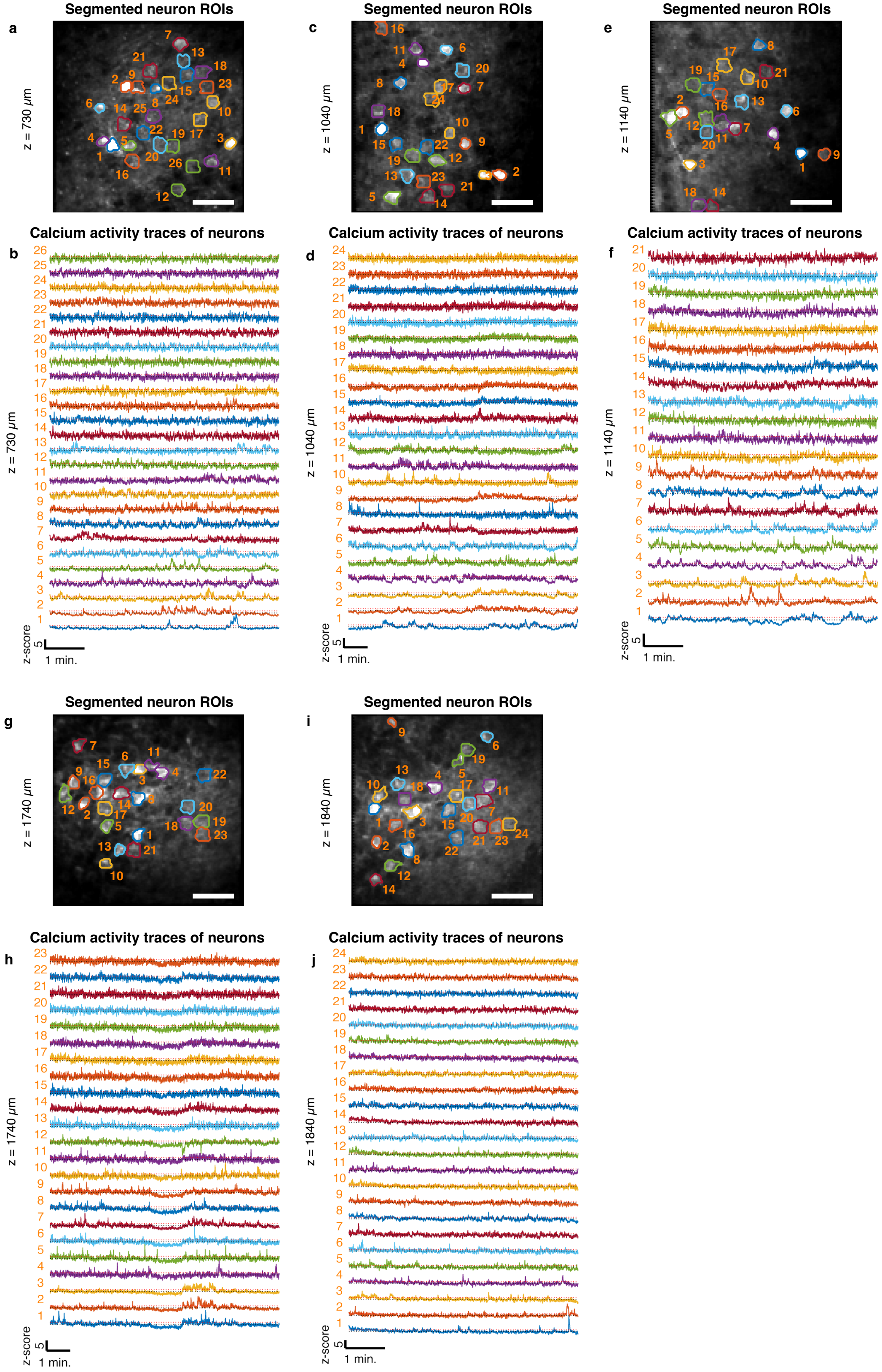

**Extended Data Fig. 6:**
**All segmented neurons and corresponding CaImAn-extracted calcium ($Ca^{2+}$) activity traces across depths in posterior parietal cortex (PPC) and hippocampus (HPC) for the data corresponding to Fig. 4.**
(**a, c, e, g, i**) Mean intensity projection of neuroactivity in individual planes covering a depth range 730–1840 µm. Segmented neurons are labeled in sequence based on their activity (z-score).
**(b, d, f, h, j**) $Ca^{2+}$ activity traces of the neurons in (a, c, e, g, i), respectively, over ~6–10 min. using CaImAn[4] with z-scores up to ~10 for the strongest responding neurons. The red dotted line denotes z-score = 1 and the black dotted line denotes the baseline.
The average power used for each of the five depths was 2.5, 6.5, 12, 58, and 66 mW, respectively. Imaging area, 230 × 230 $\mu m^2$; pixel size, 1.8 × 1.8 $\mu m^2$. Scale bar, 50 µm.

**Extended Data Fig. 7** All segmented neurons and corresponding CaImAn extracted calcium ($Ca^{2+}$) activity traces across depths within motor cortical areas and the striatum for the data corresponding to Fig. 5

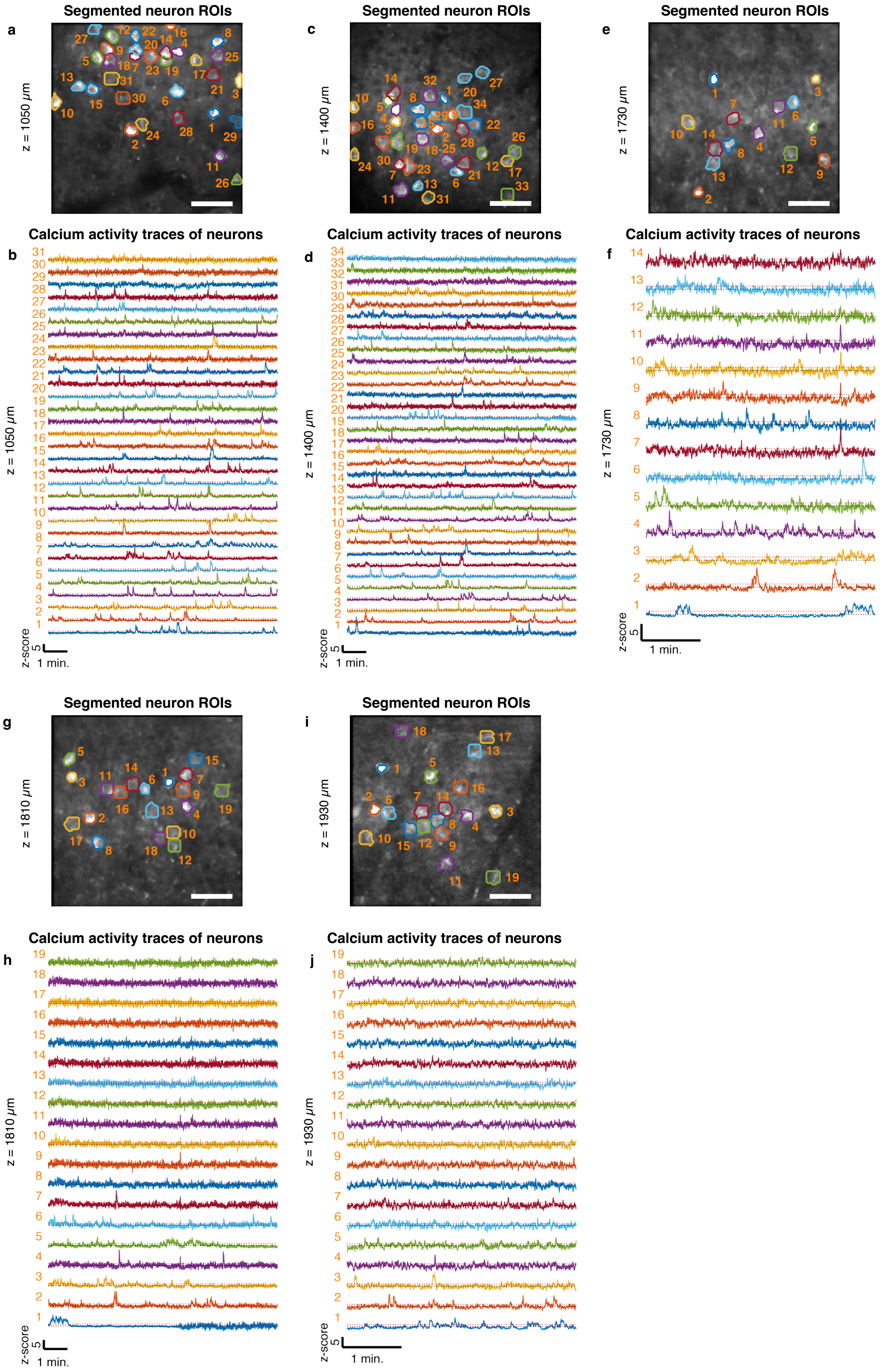

**Extended Data Fig. 7:**
**All segmented neurons and corresponding CaImAn-extracted calcium ($Ca^{2+}$) activity traces across depths within motor cortical areas and the striatum for the data corresponding to Fig. 5.**
(**a, c, e, g, i**) Mean intensity projection of neuroactivity in individual planes covering a depth range 1050–1930 µm. Segmented neurons are labeled in sequence based on their activity (z-score).
(**b, d, f, h, j**) $Ca^{2+}$ activity traces of the neurons in (a, c, e, g, i), respectively, over ~4–10 min using CaImAn[4] with z-scores up to ~12 for the strongest responding neurons. The red dotted line denotes z-score = 1 and the black dotted line denotes the baseline.
The average power used for each of the five depths was 10, 30, 120, 120, and 120 mW, respectively. Imaging area, 230 × 230 $\mu m^2$; pixel size, 1.8 × 1.8 $\mu m^2$. Scale bar, 50 µm.

#### Supplementary Video Legends

**Supplementary Video 1:**

Sequential display of individual depth planes in the mouse posterior parietal cortex (PPC) and hippocampus (HPC) brain region. Imaging FOV, 230 × 230 × 1680 $\mu m^3$; pixel size,1.8 × 1.8 × 10 $\mu m^3$. The volume stack data corresponds to the data shown in **Fig. 4b**. Scale bar, 50 µm.

**Supplementary Video 2:**

Sped-up video showing activity recordings at four different depths, data corresponding to **Fig. 4**. A rolling mean filter was applied, and the video was sped up ~ 20× for visualization. Playback was looped to match the total duration across the datasets. Imaging depths: 1040 µm (top left), 1140 µm (top right), 1740 µm (bottom left) and 1840 µm (bottom right). Imaging FOV, 230 × 230 $\mu m^2$; pixel size, 1.8 × 1.8 $\mu m^2$, Scale bar, 50 µm.

**Supplementary Video 3:**

Sped-up video showing activity recordings at four different depths, data corresponding to **Fig. 5**. A rolling mean filter was applied, and the video was sped up by~20× for visualization. Playback was looped to match the total duration across the datasets. Imaging depths: 1050 µm (top left), 1400 µm (top right), 1810 µm (bottom left) and 1930 µm (bottom right). Imaging FOV, 230 × 230 $\mu m^2$; pixel size, 1.8 × 1.8 $\mu m^2$, Scale bar, 50 µm.

#### Supplementary Files Legends

**Supplementary File 1:**

Compressed archive containing custom code for data analysis used in this work.
